# Rapid Host Plant Adaptation via Ectopic Gene Conversion in Plutellid Moths

**DOI:** 10.64898/2026.10.12.758433

**Authors:** Lisa Pormetter, Marina Pfalz, Mervic D. Kagho, Philipp Klahn, Heiko Vogel, Ute Wittstock, Juergen Kroymann

## Abstract

Herbivorous insects face an extraordinary challenge: the chemical defenses of their host plants are highly diverse, rapidly evolving, and largely unpredictable. How insects meet this challenge remains poorly understood. Here we show that ectopic gene conversion (EGC) has repeatedly driven the evolutionary diversification of glucosinolate sulfatases (GSSs), the key larval counteradaptation in plutellid moths to glucosinolate-based defenses in cruciferous plants. Across seven plutellid species, enzyme assays with heterologously expressed GSSs and 21 glucosinolates revealed distinct activity profiles: GSSs from species with broad host ranges within glucosinolate-containing plants showed balanced activity across substrates, whereas GSSs from specialists displayed profiles biased toward the glucosinolate composition of their host plants. Comparative genomic analyses identified extensive conversion among duplicated genes, both historically across the clade and in contemporary *Plutella armoraciae*. In this originally monophagous species, EGC-mediated diversification of GSS enzymes coincides with the rapid inclusion of a new cruciferous host within <200 years, demonstrating the potential of EGC to drive host-range expansion on strikingly short evolutionary timescales. Structural modelling and molecular-dynamics simulations further support functional shifts in GSS enzymes arising from EGC-derived sequence alterations. Together, these results demonstrate that EGC provides a powerful mechanism for fine-tuning detoxification enzymes, enabling herbivorous insects to adapt to chemically diverse host plants despite constrained enzymatic toolkits.

## Introduction

Plants and herbivorous insects are engaged in dynamic chemical warfare. The glucosinolate-myrosinase system, characteristic of the Brassicales, represents one of the best-characterized chemical plant defenses^1,2^. Upon tissue damage, glucosinolates are enzymatically hydrolyzed by myrosinases into reactive compounds such as isothiocyanates, which are toxic to most herbivores^3,4^ (**Fig. 1**). These defenses are highly variable across plant species, developmental stages, and environmental conditions, making them largely unpredictable to herbivores^5-8^.

**Figure 1.**
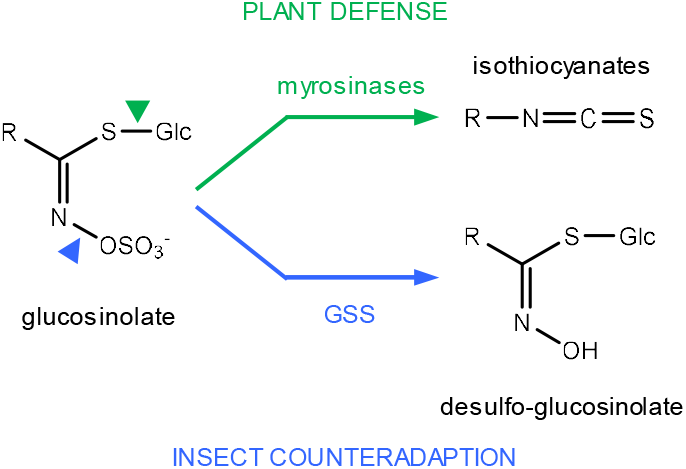
Biochemical counteradaptation in Plutellidae against the glucosinolate-myrosinase defense of Brassicales plants. Upon tissue damage, plant myrosinases activate glucosinolates, generating toxic isothiocyanates and related defense products. Larvae of Plutellidae use gut-expressed GSS to produce desulfo-glucosinolates that cannot be activated by myrosinases.

Despite the challenging complexity of glucosinolate-based defense for herbivorous insects, numerous species have specialized on glucosinolate-producing plants as their sole host plants^9,10^. For example, larvae of plutellid moths feed almost exclusively on Brassicales. Within this group, different species exhibit distinct host plant preferences and host ranges – from narrow specialists to more flexible feeders. Among them, the polyphagous *Plutella xylostella* is recognized as the most destructive global pest of cruciferous crops such as cabbage, broccoli, and canola^11^. Larvae of *P. xylostella* possess glucosinolate sulfatases (GSS), enzymes that remove the sulfate moiety from glucosinolates, converting them into desulfo-glucosinolates^12,13^. This transformation renders the compounds inert to myrosinase, effectively neutralizing the plant defense mechanism before it can be triggered (**Fig. 1**). Other plutellid species may use the same mechanism to thrive on glucosinolate-containing plants; in this case, variation in GSS activity and substrate specificity could explain their differential host use.

The extraordinary diversification of plant chemical defenses places substantial demands on herbivore detoxification systems^14,15^. For example, the enzymes encoded by three *GSS* genes in *P. xylostella* must cope with the vast diversity of glucosinolate profiles produced across the exceptionally broad host range of this species^16,17^. Even in oligo- or monophagous plutellids, individuals feeding across different plants within a population – or shifting among populations or habitats – encounter host glucosinolate chemistries that may be novel and potentially unmanageable^8^. Such a persistent mismatch between plant chemical innovation and insect metabolic capacity is expected to impose strong selection for detoxification flexibility, favoring mechanisms such as substrate promiscuity, context-dependent gene expression, and gene family remodeling.

Ectopic gene conversion (EGC) is a non-reciprocal transfer of sequence information between paralogous genes, producing homogenized or mosaic alleles through aberrant recombination during DNA repair^18,19^. By introducing multiple coordinated amino-acid changes in a single event, EGC can modify protein function more rapidly than point mutation. EGC has long been hypothesized to facilitate functional innovation in gene families^20,21^, but direct evidence for ecologically meaningful consequences remains scarce. Here, we test the role of EGC in shaping the functional diversification of detoxification enzymes and its contribution to host-plant adaptation, as exemplified by the GSS of plutellid moths.

## Results

### *GSS* monophyly in the Plutellidae

Throughout the manuscript, we use abbreviations for glucosinolates (Extended Data Table 1) and group them by side-chain structure: alkyl glucosinolates: 1-me, 2-mp, 1-mp; alkenyl glucosinolates: 2-prop, 3-but, 4-pent; methylthioalkyl glucosinolates: 4-mtb; methylsulfinylalkyl glucosinolates: 3-msop, 4-msob, 5-msop, 6-msoh, 8-msoo, 10-msod; methylsulfonylalkyl glucosinolates: 3-msoop; benzenic glucosinolates: 2-pe, pohb, benzyl; indole glucosinolates: i3m, 4ohi3m, 4msoi3m, 1moi3m.

To compile an inventory of *GSS* genes in representative species of the Plutellidae for phylogenetic analyses, we generated near-complete genome assemblies from single specimens of *Plutella armoraciae* (*Par*), *Plutella porrectella* (*Ppo*), *Eidophasia messingiella* (*Eme*), *Leuroperna sera* (*Lse*) and *Rhigognostis hufnagelii* (*Rhu*), along with the outgroup *Baerenschenkia umtrunkula* (Extended Data Table 2). Haploid genome sizes ranged from 349 to 989 Mbp (Extended Data Table 3). Together with *P. xylostella*^22,23^ (*Pxy*) and *P. australiana*^24^ (*Pau*), this provided genomic data for seven plutellid species.

All near-complete plutellid genomes contained at least one homolog each of *GSS1* and *GSS3*, whereas *B. umtrunkula* lacked these genes. *GSS1* copy number ranged from one in *P. australiana* and *R. hufnagelii* to four in *E. messingiella*, and only *R. hufnagelii* carried an additional *GSS3* paralog (Extended Data Table 4). We frequently recovered both alleles per gene.

Gene trees of *GSS*, inferred using Bayesian methods, strongly supported a single, monophyletic origin for all *GSS* genes within the Plutellidae. However, the branching topologies of the *GSS1* and *GSS3* subtrees exhibited marked incongruence (**Fig. 2**). The basal branch position was occupied by *L. sera* in the *GSS1* subtree but by *R. hufnagelii* in the *GSS3* subtree. Furthermore, *GSS1* genes of *E. messingiella* formed a distinct clade, separate from the monophyletic group comprising all *GSS1* genes of the *Plutella* genus. In contrast, the *GSS3* gene of *E. messingiella* clustered with those of *P. armoraciae* and *P. porrectella*, forming the sister clade to *P. xylostella* and *P. australiana*. Finally, *ParGSS1-1a2* clustered robustly alongside *ParGSS1-2*, despite occupying the same genomic position upstream of a *ParGSS1-2* allele as *ParGSS1-1a1* (Extended Data Fig. 1).

**Figure 2.**
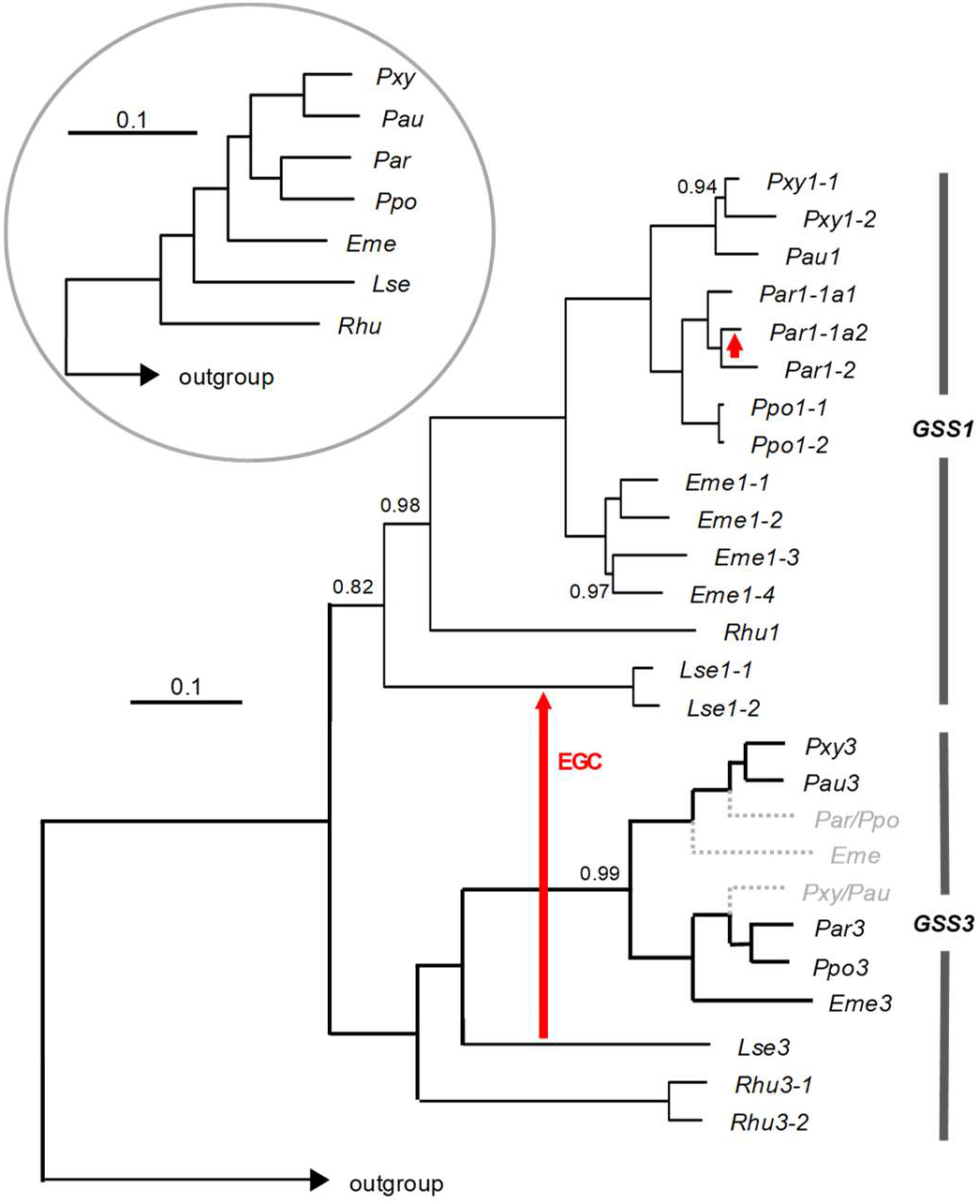
*GSS* gene tree and mitochondrial phylogeny. Bayesian gene tree of *GSS* sequences (posterior probability of 74.2%) and corresponding mitochondrial phylogeny (insert; posterior probability 100%). Scale bars indicate the expected number of substitutions per codon site. Except for nodes marked by juxtaposed numbers, all nodes in both trees received full posterior support. Dotted gray lines denote hypothetical deletions of *GSS3* genes following a cryptic duplication. Red arrows indicate inferred ectopic gene conversion events, including sequence transfer from *Leuroperna sera GSS3* to *GSS1* and conversion between *GSS1* paralogs in *Plutella armoraciae*.

To explore the causes of these inconsistencies, we first inferred a phylogeny from concatenated mitochondrial protein-coding genes. The resulting, strongly supported topology placed *R. hufnagelii* as the basal branch, followed by *L. sera, E. messingiella*, and the four *Plutella* species. We recovered the same topology for *SulfD*, a gene closely linked to the *GSS* cluster^13^ (Extended Data Fig. 2).

Next, we compared sequence identities of the *GSS1* and *GSS3* coding sequences across species (Extended Data Table 5). Sequences were 0.9 to 4.7 percentage points more similar within species than between species (*t* ≥ 5.1, *df* = 6, *p* ≤ 0.01), indicating non-independent paralog evolution. Together with the phylogenies, this pattern strongly supported ECG as a driver of GSS gene family evolution.

### Evidence for EGC

To test whether EGC could account for our observations, we developed a novel statistical framework, the ‘OTTO-TOTO test’. After gene duplication and subsequent speciation, orthologs – gene copies in different species that originated from a common ancestral gene – are typically more similar to one another than to paralogs, which are duplicated genes within the same genome. The acronyms OTTO and TOTO describe alternative patterns of nucleotide identity at individual alignment sites across the four sequences representing gene copies 1 and 2 in species A and B. In the OTTO configuration, at a given site, copy 1 in species A shares its nucleotide with copy 2 in species B, while copy 2 in A shares a different nucleotide with copy 1 of B (Extended Data Fig. 3). In contrast, the TOTO configuration arises when, at that same site, the two paralogs in species A share one nucleotide and the two paralogs in species B share a different one. Under independent paralog evolution, OTTO and TOTO configurations should occur with similar frequency. A significant excess of TOTO over OTTO instead indicates non-independent paralog evolution, consistent with EGC.

Across all pairwise species comparisons of *GSS1* and *GSS3*, OTTO and TOTO configurations increased with phylogenetic distance, with TOTO accumulating substantially faster than OTTO (Extended Data Fig. 4). This pattern resulted in a consistent excess of TOTO over OTTO configurations across species, reflecting pervasive non-independent paralog evolution.

Next, we used TOTO-compatible sites to map the phylogenetic distribution of EGC, assess whether these conversions altered amino-acid sequence, and, where possible, infer their direction. EGC occurred on every major branch, including the most recently diverged lineages, and most species showed evidence for bidirectional conversion between *GSS1* and *GSS3* (Extended Data Fig. 5; Extended Data Table 6). Of the 286 sites compatible with EGC, only 84-90 were nonsynonymous, indicating that EGC is generally constrained by purifying selection, consistent with previous findings^25^. A notable exception occurred in *L. sera*, where 26-29 of 54 inferred changes were nonsynonymous. Here, directional inference revealed only a single case in which *GSS1* replaced *GSS3* sequence, whereas 17 cases involved replacement of *GSS1* sequence with *GSS3*-derived nucleotides – indicating a strong, asymmetric invasion of the *GSS1* locus by *GSS3*. Consequently, *L. sera GSS1* genes appeared unusually similar to *GSS3*, causing the *L. sera GSS1* sequences to cluster artificially near the base of the *GSS1* subtree.

Pairwise comparisons of genomic *ParGSS1* sequences showed that *ParGSS1-1a1* and *ParGSS1-1a2* shared identical intron structures without major indels, and *ParGSS1-2a1* and *ParGSS1-2a2* likewise differed only by a few, small indels (Extended Data Fig. 6). In contrast, comparisons between either *ParGSS1-1* allele and *ParGSS1-2a1* revealed numerous indels, consistent with the two *ParGSS1-1* copies forming a distinct allelic pair. Notably, however, intron 3 was markedly more similar between *ParGSS1-1a2* and *ParGSS1-2a1*, indicative of EGC.

To identify the converted regions, we examined synonymous and non-synonymous polymorphisms distinguishing *ParGSS1-1a2* and *ParGSS1-2a1* from *ParGSS1-1a1*. Several positions were shared exclusively between *ParGSS1-1a2* and its paralog *ParGSS1-2a1* (Extended Data Fig. 6), clustering in exons 4 and 7. This pattern indicates at least two EGC events in which segments of *ParGSS1-1a2* were replaced by *ParGSS1-2* sequence, causing the artificial clustering of these paralogs in the gene tree.

By contrast, the divergent placement of *E. messingiella* in the *GSS1* and *GSS3* subtrees was best explained by a cryptic duplication of the *GSS3* gene in the common ancestor of *Eidophasia* and *Plutella*, followed by reciprocal loss: one copy was lost in *Eidophasia* and in the shared ancestor of *P. armoraciae* and *P. porrectella*, whereas the alternative paralog was lost in the lineage leading to *P. xylostella* and *P. australiana*.

Early analyses using standard tests for positive selection with PAML^26^ and aBSREL^27^ suggested widespread adaptive evolution across *GSS1* and *GSS3* (Extended Data Fig. 7). However, these methods assume that substitutions arise independently through point mutations, an assumption violated by the extensive EGC documented here. Consequently, signals of positive selection cannot be interpreted independently of EGC and are not considered further.

### Functional variation of GSS enzymes

We expressed the corresponding cDNAs, along with three outgroup cDNAs, in *Sf*9 cells to determine whether *GSS* genes encode functional enzymes with activity on structurally diverse glucosinolates. Culture supernatants were buffer-exchanged and incubated with an equimolar mixture of 21 glucosinolates (Extended Data Table 1). After 14 h, UHPLC-HRMS analysis detected desulfo-glucosinolates in reactions from 27 of 29 tested enzymes but not in negative controls, demonstrating GSS activity (Extended Data Table 7). Two inactive reactions likely reflected insufficient expression, as supported by Western blot analysis.

Of the 27 enzymes that showed activity, 25 classified as GSS based on phylogenetic analyses were examined in greater detail. After 14 h of incubation, the majority of these enzymes achieved near-complete conversion of the 21 glucosinolates, confirming their ability to act on a wide structural range. Shorter incubations, however, revealed pronounced differences between GSS1- and GSS3-type enzymes (**Fig. 3**). GSS1 enzymes preferentially converted aliphatic and benzenic glucosinolates, and showed weak activity toward indole glucosinolates. Among aliphatic glucosinolates, their activity varied across branched chain amino acid (BCAA)-derived and methionine (Met)-derived compounds.

**Figure 3.**
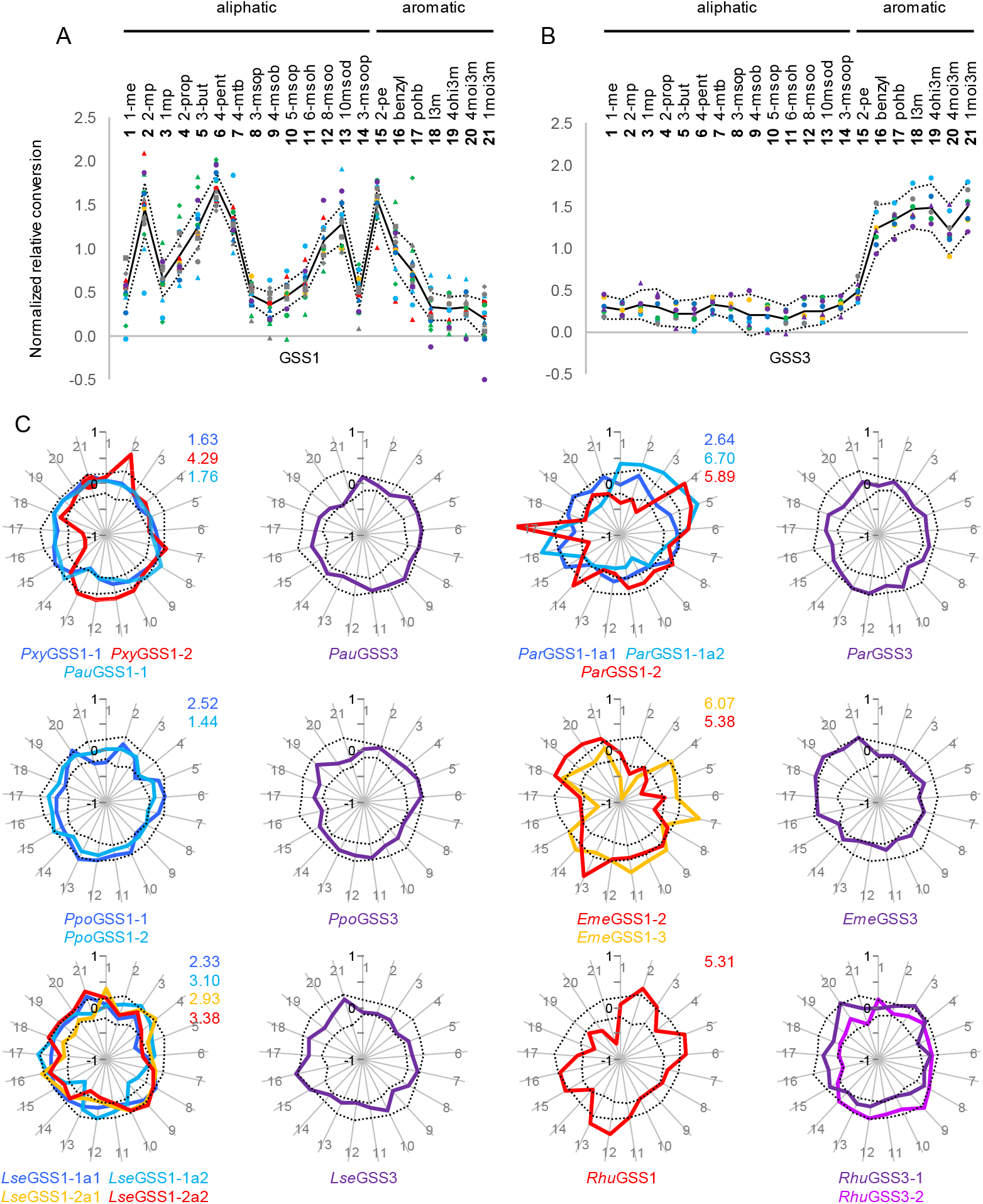
Enzymatic footprints of GSS1 and GSS3 in Plutellid moths. **A, B.** Normalized relative conversion of 21 glucosinolates by GSS1-type (**A**) and GSS3-type (**B**) enzymes (means of three expressions). Each colored symbol represents one enzyme; solid lines indicate global means across all GSS1 (N = 15) or GSS3 (N = 7), with 99 % confidence intervals shown as dotted lines. **C**. Differential enzymatic footprints represented as spider charts. Each axis corresponds to one glucosinolate (numbering as in panels A and B), with the global mean of its normalized relative conversion set to zero. Colored lines show deviations of individual enzymes from these means; different colors denote different enzymes within a species. *P. xylostella* and *P. australiana* GSS1 are combined in one chart. Colored numbers in the upper right corner of GSS1 panels indicate the corresponding deviation indices (sum of absolute deviations across all substrates).

In contrast, GSS3 enzymes displayed limited activity toward aliphatic glucosinolates and the benzenic 2-pe, but readily converted other aromatic glucosinolates, especially indole glucosinolates.

Correlation analysis of GSS1 activities across 21 substrates resolved four major substrate clusters (Extended Data Fig. 8). BCAA-derived glucosinolates together with benzyl glucosinolate formed an intercorrelated cluster that showed only weak or mixed correlations with alkenyl substrates but negative correlations with methylsulfinylalkyl and indole glucosinolates. Alkenyl glucosinolates formed a second cluster with moderate internal correlations and negative correlations with short-chain methylsulfinylalkyl and indole substrates. Oxidized Met-derived glucosinolates (methylsulfinylalkyl and methylsulfonylalkyl) formed a distinct cluster in which correlation strength depended on side-chain length: substrates with similar chain lengths were highly intercorrelated, whereas correlations between short- and long-chain derivatives were weak. Oxidized Met-derived glucosinolates also showed negative correlations with benzyl and 2-pe. Indole glucosinolates constituted a fourth, strongly intercorrelated cluster characterized by pronounced negative correlations with alkenyl substrates and weak positive correlations with medium-chain methylsulfinylalkyl glucosinolates. Activity toward 2-pe did not correlate significantly with any other activity.

Correlation analysis of GSS3 activities resolved a strongly bipartite structure (Extended Data Fig. 8). Aliphatic substrates formed a single, broadly intercorrelated cluster. Aromatic glucosinolates formed a second, tightly coherent cluster, although correlations between 2-pe and the other aromatic glucosinolates were only weak. Correlations between the aliphatic and the aromatic cluster were generally negative, reflecting a pronounced aliphatic-aromatic separation in GSS3 substrate preferences, consistent with previous findings^13^.

To visualize differential activity patterns, we normalized each enzyme’s activity toward each substrate by its total activity across all substrates and calculated substrate-specific global means with 99% confidence intervals, averaging across all GSS1 or all GSS3 enzymes for each substrate. Activity patterns showed that both GSS1- and GSS3-type enzymes exhibited narrow activity ranges across substrates, consistent with functional constraints on substrate use (**Fig. 3A, B**). Deviation indices, defined as the sum of absolute deviations from the global mean (**Fig. 3C**), revealed three principal classes of GSS1 enzymes. The first class comprised enzymes with low deviation indices and uniform, low-bias activity profiles (*Pxy*GSS1-1, *Pau*GSS1-1, and *Ppo*GSS1-2). The second class consisted of enzymes with a pronounced substrate bias, characterized by high deviation indices (*Pxy*GSS1-2, *Par*GSS1-2a2, *Par*GSS1-2, *Eme*GSS1-2, *Eme*GSS1-3, and *Rhu*GSS1). The third class displayed intermediate profiles (*Par*GSS1-1a1, *Ppo*GSS1-1 and all four *L. sera* variants). Notably, several *Par*GSS showed pronounced deviations from the global means specifically for 1-me and 2-prop, motivating their inclusion in subsequent modeling analyses.

Whenever multiple GSS1 paralogs were present, they exhibited divergent activity profiles, and allelic variants differed likewise. In contrast, GSS3 enzymes showed rather homogeneous activity profiles, with *Eme*GSS3 showing the lowest relative activity toward aliphatic and *Rhu*GSS3-1 vs. *Rhu*GSS3-2 exhibiting slightly complementary profiles (**Fig. 3C**).

To assess activity toward unusual substrates, we incubated GSS1 and GSS3 from *P. australiana* and *P. porrectella* with glucosinolate extracts from *Hesperis matronalis*, which produces apiosylated benzenic glucosinolates^8,28^. GSS1 enzymes from both species showed virtually no activity toward these specific glucosinolates, but readily converted 4-pent and benzyl glucosinolate added to the extract (Extended Data Fig. 9). In contrast, both *Pau*GSS3 and *Ppo*GSS3 processed apiosylated glucosinolates.

### Enzyme-substrate interactions

To assess how EGC affects performance of the encoded enzymes, we modelled *Par*GSS1-1a1, *Par*GSS1-1a2, and *Par*GSS1-2 in complex with 1-me and 2-prop (**Fig. 4**). We also included variants of *Par*GSS1-1a1 carrying the ectopically converted residues in exon 4 (1-1a1_e4), exon 7 (1-1a1_e7), or both regions (1-1a1_e4e7). These substitutions did not alter the architecture of the binding pocket relative to wild-type *Par*GSS1-1a1.

**Figure 4.**
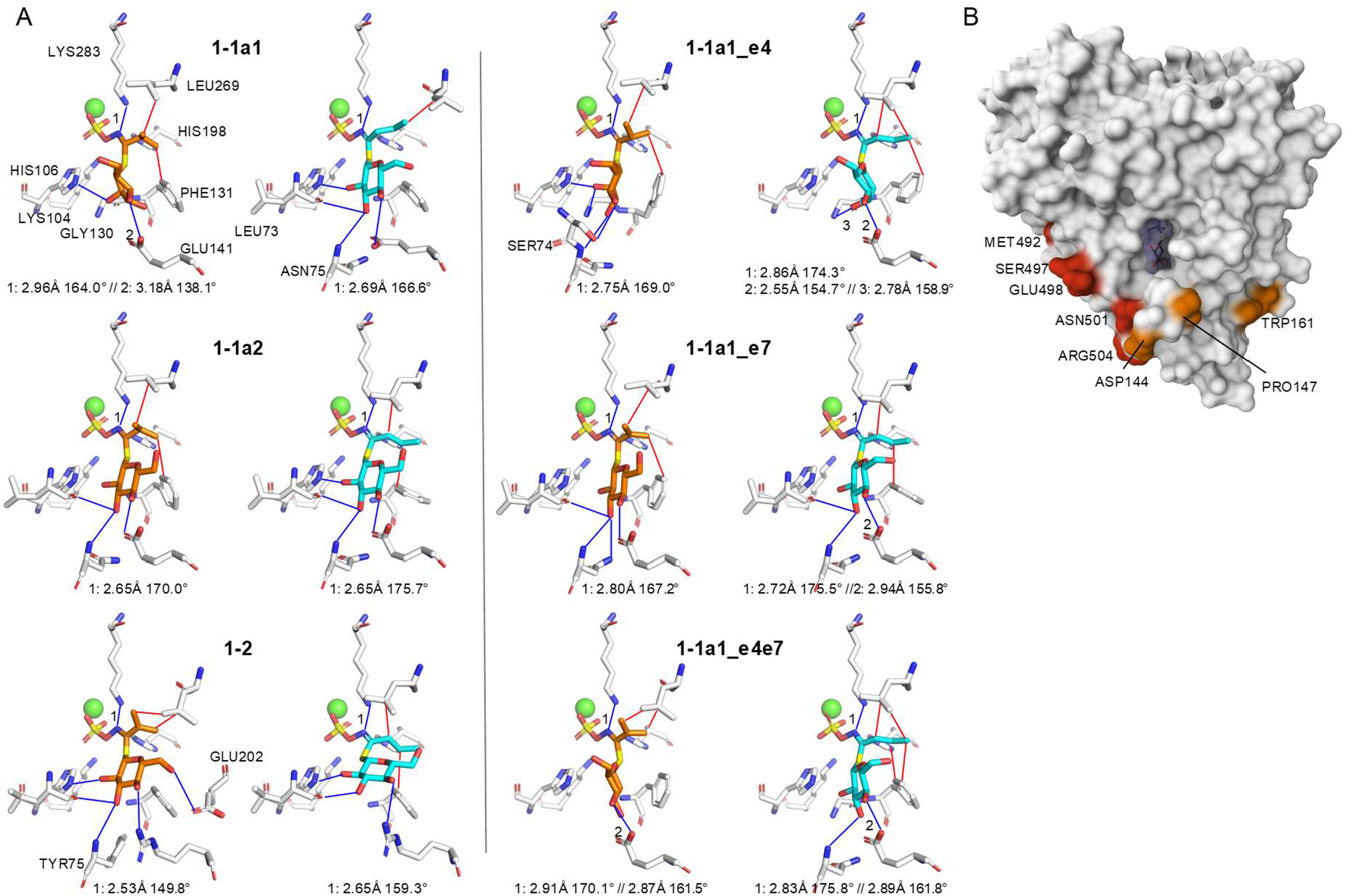
Ligand binding modes of *P. armoraciae* GSS1 enzymes and EGC variants. **A**. Predicted binding poses of 1-me (orange) and 2-prop (cyan) for *Par*GSS1-1a1, *Par*GSS1-1a2 and *Par*GSS1-2, and for *Par*GSS1-1a1 variants derived after exon-specific replacements of ectopically converted residues (exon 4, exon 7 or both). Ca^2+^ is shown in green; PLIP-predicted hydrogen bonds in blue; hydrophobic contacts in red. The canonical arylsulfatase binding architecture – including the catalytic *α*-formylglycine and conserved Ca^2+^-coordinating and sulfate-binding residues^30,31^ – is omitted for clarity. The Ca^2+^-sulfate ester geometry remains conserved; and the glucosinolate adopts the same principal orientation across enzymes. Hydrogen bond values correspond to donor-acceptor distances ≤ 3.3 Å and donor-hydrogen-acceptor angles ≥ 130°. **B**. Surface representation of *Par*GSS1-1a2 with bound 1-me. Ectopically converted residues are highlighted in orange (exon 4) and red (exon 7). Thr176, Ile178 and Arg491 are not visible in this orientation; only Ile178 is not surface-exposed.

Structural confidence metrics (Extended Data Table 8) were uniformly high across all modelled enzymes, indicating stable folds and well-defined active-site architectures. *Par*GSS1-1a2 showed slightly higher structural confidence than *Par*GSS1-1a1 or *Par*GSS1-2. Introducing the converted residues from *Par*GSS1-1a2 into *Par*GSS1-1a1 increased overall structural confidence, with both exon 4 and exon 7 contributing.

As the best-scoring models exhibited similar binding poses across enzymes (**Fig. 4**), we performed molecular-dynamics simulations with 1-me to assess substrate delivery under conformational flexibility. The ligand was placed ∼2 nm from the Ca^2+^ ion in the active center, and the system was equilibrated to avoid immediate collapse. We simulated 20 ns from ten randomly generated starting poses per enzyme, with two replicates each (400 ns per enzyme). Throughout simulations, we monitored the sulfate-ester-to-Ca^2+^ distance (*d*).

Across enzymes, approximately half of the trajectories showed ligand escape (*d* > 3.6 nm). Among trajectories remaining within this threshold (*d* ≤ 3.6 nm), residency times did not differ significantly (*F*_5,112_ = 1.33; *p* = 0.257; *N*=19-20; Extended Data Table 9), indicating that differences arose primarily from how efficiently each enzyme delivered the ligand into the near-contact region.

Distance distributions for the three wild-type enzymes revealed pronounced differences in near-contact delivery (**Fig. 5**). *Par*GSS1-1a1 rarely sampled short distances (*d* < 0.7 nm), whereas *Par*GSS1-1a2 showed a strong peak at 0.43-0.55 nm, adjacent to the catalytically competent pose (*d* ≈ 0.3 nm). *Par*GSS1-2 displayed an intermediate pattern.

**Figure 5.**
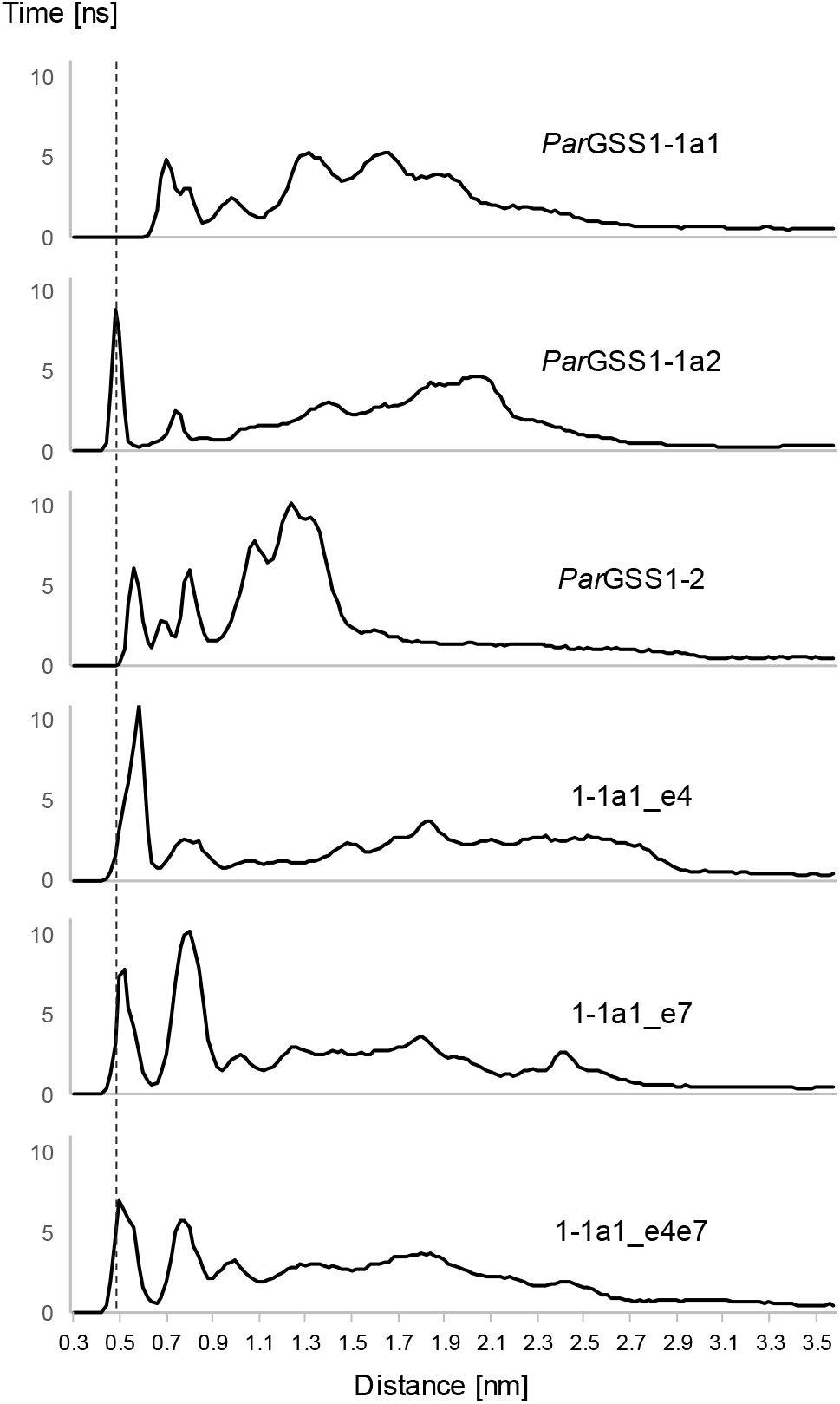
Residency-based quantification of substrate proximity. Distances between the sulfate ester group of 1-me and the Ca^2+^ in the catalytic center were extracted from MD trajectories and binned into 0.2-nm intervals. Frame counts per bin were summed across trajectories and converted into residency time (ns). Values > 3.6 nm were excluded. The vertical dotted line marks the peak position of *Par*GSS1-1a2. Distances around 0.3 nm correspond to the bound configuration defined by the static structural models, whereas distances > 3.6 nm indicate loss into bulk solvent.

All three conversion variants improved near-contact delivery relative to *Par*GSS1-1a1 and *Par*GSS1-2. Variants carrying exon 7 (1-1a1_e7 and 1-1a1_e4e7) closely resembled *Par*GSS1-1a2, with high occupancy at *d* < 1.0 nm and clear enrichment in the 0.4-0.6 nm interval. Exon 4 conversion alone produced an intermediate phenotype. Together, these results are consistent with EGC introducing sequence changes that enhance ligand delivery into the catalytically competent region.

## Discussion

### Insect adaptation to host glucosinolates

GSS1 and GSS3 both act on a wide range of glucosinolates, and – given sufficient time – can desulfate all major glucosinolate classes. This inherent breadth is illustrated by GSS3 from *P. australiana*, which metabolizes the major apiosylated glucosinolate from *Hesperis matronalis*, a plant species the insect never encounters in its native habitat. However, larval development is time-critical, and delays markedly increase mortality^32-34^. Thus, despite this broad catalytic potential, selection favors GSS activity patterns that match the glucosinolate profiles of an insect’s preferred hosts.

Polyphagous insects with broad host ranges encounter diverse und unpredictable glucosinolate mixtures. Consequently, selection favors GSS1 activity patterns that ensure reliable performance across substrates rather than specialization toward individual glucosinolates. In *P. xylostella*, this generalist pattern is reinforced by high constitutive expression of *GSS1* genes, and inducible expression of *GSS3* for indole glucosinolates^13^. *P. australiana*, which diverged from *P. xylostella* ∼2 million years ago^24^, retains a GSS1 enzyme with an almost identical substrate spectrum, suggesting that this generalist activity has been conserved in both species. However, the second *P. xylostella* GSS1 enzyme, *Pxy*GSS1-2 shows hallmarks of specialization, with a pronounced bias toward medium-to long-chain Met-derived glucosinolates as well as 2-mp. *Pxy*GSS1-2 enhances detoxification efficiency on hosts enriched in these glucosinolates and complements the broadly active *Pxy*GSS1-1 enzyme by reducing substrate competition across glucosinolate classes.

The third polyphagous species in our sample, *L. sera*, currently feeds on a broad range of wild and cultivated Brassicaceae. Yet, its GSS activity profiles suggest that this generalist state is evolutionary recent and not yet fully consolidated. The two GSS1 enzymes, *Lse*GSS1-1 and *Lse*GSS1-2, display complementary activity profiles, and all allelic variants examined show intermediate deviation indices with widely distributed – though not yet stabilized – generalist activity across substrates. Strikingly, these biochemical signatures coincide with the species’ ongoing geographic expansion from its East Asian origin to much of Asia and Oceania^35,36^.

By contrast, insects with restricted host ranges – such as *E. messengiella, R. hufnagelii* or *P. armoraciae* − tend to exhibit GSS1 activity patterns biased towards the glucosinolate profiles of their host plants. For example, in *R. hufnagelii*, the single GSS1 enzyme shows above-average relative activity toward BCAA-derived, long-chain Met-derived, and alkenyl glucosinolates, as well as 2-pe – substrates that closely match the glucosinolate composition of its host *Arabis hirsuta*^37-39^.

The most striking example of host-aligned specialization, however, is *Plutella armoraciae*. Native to North America, it has the most restricted host range of the species examined, feeding primarily on horseradish (*Armoracia rusticana*). Horseradish foliage contains exceptionally high glucosinolate concentrations, with 2-prop typically accounting for ∼90% of the total glucosinolate content. In addition, around a dozen aliphatic, benzenic, and indole glucosinolates are present, though their relative abundance varies by cultivar, plant age, and environmental conditions; BCAA-derived glucosinolates, if present, occur only at trace levels^40-42^. This compositional profile is mirrored in the activities of *Par*GSS1-1a1, *Par*GSS1-2, and *Par*GSS3 (**Fig. 3B**): *Par*GSS1-2 is specialized for alkenyl glucosinolates, particularly 2-prop; *Par*GSS1-1a1 likely acts on minor aliphatic glucosinolates, and *Par*GSS3 targets aromatic glucosinolates.

In southwestern Canada, *Sisymbrium loeselii* serves as an alternative host for *P. armoraciae*^43^. Introduced to North America in the 1850s, the earliest documented record of *S. loeselii* on the continent in the Global Biodiversity Information Facility dates to 1871 (https://doi.org/10.15468/dl.jaa9k5), and the species has since spread widely across the continent. *S. loeselii* is characterized by two BCAA-derived glucosinolates, 1-me and 1-mp, in addition to indole glucosinolates^44^. Notably, 1-me accumulates in high concentrations in the shoots, reaching levels comparable to those of 2-prop in horseradish leaves. On this new host, *Par*GSS1-1a2 plays a key role, as this enzyme is specialized for BCAA-derived glucosinolates alongside alkenyl glucosinolates (**Fig. 3B**).

### EGC as an accelerator of adaptation

The rapid inclusion of *S. loeselii* into *P. armoraciae*’s host range – within <200 years of its introduction − coincides with biochemical signatures of adaption in GSS enzymes. Our genomic and functional data identify EGC as a key mechanism underlying this adaptation, enabling the transfer of multiple substitutions that enhance 1-me desulfation.

*GSS1* and *GSS3* are more similar within than between species (Extended Data Table 5), and OTTO-TOTO tests revealed an excess of polymorphisms consistent with EGC (Extended Data Fig. 4). Across the phylogeny, signatures of EGC are frequent, and in some lineages – most notably *L. sera* – conversion has proceeded directionally, replacing segments of *GSS1* with *GSS3*-derived sequences. These patterns indicate that EGC has repeatedly reshaped *GSS* evolution over long timescales, occasionally with strong directional bias.

The OTTO–TOTO test is informative when gene duplication predates speciation events; however, all *GSS1* duplications appear to be species-specific. Nonetheless, while contemporary *GSS1* and *GSS3* coding sequences share just over 70% identity, even young branches of the phylogeny show clear evidence of EGC between *GSS1* and *GSS3* (Extended Data Figs. 4, 5). Because EGC frequency increases with sequence similarity^45,46^, conversion among contemporary *GSS1* paralogs is likely common. Recurrent homogenization between *GSS1* copies may thus explain why all duplications within the *GSS1* subtree appear to be species-specific.

In *P. armoraciae, ParGSS1-1a2* and *ParGSS1-2a1* share multiple identical nucleotides at positions where *ParGSS1-1a1* carries different bases, including nonsynonymous changes clustered in exons 4 and 7, consistent with at least two conversion events (Extended Data Fig. 6). Although these changes do not directly alter the binding pocket or catalytic center (**Fig. 4**), they nonetheless enhance acceptance of 1-me – the major novel glucosinolate encountered on *S. loeselii* – by *Par*GSS1-1a2. Molecular dynamics simulations show that introducing these converted residues into *Par*GSS1-1a1 shifts the substrate trajectory closer to the active center (**Fig. 5**), providing a molecular basis for this enhanced specificity. Together, these results strongly support EGC as a mechanistic driver of rapid adaptation in *P. armoraciae* to its new host plant.

Gene duplication is pervasive across genomes^47-49^, providing abundant substrates for EGC. Although duplicated genes tend to functionally diverge over time, ongoing EGC often homogenizes paralogs^20^. Consequently, strong selection is required to establish and maintain divergent copies. Critically, EGC does not introduce random mutations; instead, it transfers amino acid substitutions already functionally validated in the donor paralog. Thus, recipient copies rarely lose activity entirely. Rather, such transfers typically recalibrate enzyme function, shifting substrate preference or modulating catalytic properties, as exemplified by the GSS enzymes of *Plutella armoraciae*. Under stable environments, this redistribution can disrupt optimized combinations of paralogs and lead to maladaptation^25^. However, in changing environments, EGC can deliver variants that accelerate adaptation relative to point mutations alone. Hence, this dual potential to either erode or facilitate adaptation depends on ecological context, and may enable rapid responses to contemporary environmental change, including climate-driven shifts in host plant availability or chemistry.

## Methods

### Source material and sequencing

Insect specimens were generously provided by micro-moth experts. High molecular weight (HMW) DNA was extracted from individual larvae or adult specimens using the Genomic tip kit, and DNA concentrations were measured with a Qubit 4 Fluorometer using the Qubit 1x sdDNA Assay. To enrich for long DNA fragments in samples with higher fragmentation levels, the Short Read Eliminator kit was applied, and remaining impurities were removed using AMPure XP beads. DNA quality and size distribution were assessed on a TapeStation System. For PacBio HiFi sequencing^50^, library preparation and HiFi sequencing were performed by the Max-Planck Genome Center, Cologne, on a PacBio Sequel II instrument. Genome assemblies were generated using hifiasm v0.16.1^51^. For Oxford Nanopore Technologies (ONT) sequencing^52^, end-DNA repair was performed using the NEBNext Ultra II DNA Library Prep Kit, followed by ligation of sequencing adapters from the V14 Ligation Sequencing Kit. Sequencing was carried out on a MinION platform using R10.4.1 flow cells. Super high-accuracy (SUP) base-calling was performed with Dorado v1.3.2 (v5.2 DNA models) implemented in MinKnow. ONT genome assemblies were generated using Flye v2.9.2^53^, followed by four rounds of polishing with Racon and one round of error correction with Medaka. Final polishing was performed using ntEdit with paired-end (2x 150bp) Illumina reads^54^ generated from the same HMW DNA. PurgeDups was used to remove duplicated (heterozygous) regions and generate haploid assemblies. Taxon IDs were assigned to contigs using BlobTools v3.3.4 and TaxIdent^55^, and bacterial, fungal and plant contaminants were removed. Genome completeness was assessed using BUSCO v5.3.0 with the Insecta dataset (1367 single-copy orthologs) from OrthoDB v10.

### Sequence alignment and phylogenetic inference

Multiple sequence alignments were generated with MUSCLE5^56^ via the EMBL-EBI platform. The *GSS* alignment comprised 496 codons, excluding signal-peptide codons and gap positions; the *SulfD* alignment comprised 504 codons; and the concatenated mitochondrial protein-coding genes comprised 3708 codons. Phylogenetic analyses were performed in MrBayes v3.2.7^57^ using a codon-based substitution model under the general time reversible framework, with site-specific variation in nonsynonymous-to-synonymous substitution rate ratios modeled under the M3 discrete distribution. Six Markov chains were run for 500,000 (*GSS, SulfD*) or 100,000 (mitochondria) generations, allowing two swaps between chains to enhance mixing, and sampling every 2500 or 500 generations, respectively. The first 25% of samples were discarded as burn-in, and majority-rule consensus trees were constructed with posterior probabilities estimated for each clade. Outgroups were *Yponomeuta cagnagella* and *Baerenschenkia umtrunkula* for *GSS, Y. cagnagella* for *SulfD, and Ypsolopha scabrella* and *B. umtrunkula* for the mitochondrial phylogeny.

### Dot-plot and ancestral reconstruction analyses

Dot-plot analyses were performed using YASS^58^, and ancestral sequence reconstruction was carried out with FastML^59^.

### Comparison of GSS1 and GSS3 coding sequences

Pairwise sequence identities were calculated in DNASTAR from codon-based alignments. For each species, mean identity between its *GSS1* and *GSS3* genes and alleles was computed. A second mean was calculated between the same *GSS1* sequences and all *GSS3* sequences from other species. Differences between within-species and between-species identities were assessed using a two-sided paired *t*-test implemented in jamovi^60^).

### Heterologous expression

cDNAs encoding the arylsulfatase-like genes of *P. xylostella* (*PxyGSS1-1, PxyGSS1-2, PxyGSS3, PxySulfD*) were amplified by PCR using gene-specific primers containing a 5′ Kozak sequence. Amplicons were ligated into the pIB/V5-His TOPO-TA vector (Invitrogen) as described^13^. For the remaining Plutellidae *GSS* genes, codon-optimized open reading frames with a 5′ Kozak sequence were synthesized (Genscript) and cloned in the pIZ/V5-His vector (Invitrogen) via *Hind*III/*Sac*II restriction sites. All constructs were verified by Sanger sequencing for correct orientation and insert identity. Native stop codons were omitted to enable C-terminal fusion to a V5-(His)6 epitope. *Sf*9 cells were maintained at 27 °C in *Sf*-900 II SFM and transfected with FuGENE HD according to the manufacturer’s instructions. After 72 h, culture supernatants were cleared by centrifugation (4000 × g, 10 min, 4 °C) and concentrated with buffer exchange in 100 mM Tris-HCl (pH 7.5). A non-transfected control was processed identically. Recombinant protein levels were estimated by dot blot using anti-V5-HRP and quantified in ImageJ^61^ relative to a calibration standard; samples were diluted to equal enzyme concentrations.

### GSS assay with glucosinolate mix

A mixture of 21 glucosinolates (25 µl; Extended Data Table 1) was combined with 25 µl enzyme solution in a final volume of 100 µl 50 mM Tris-HCl (pH 7.5), yielding 25 µM of each glucosinolate. Negative controls used boiled enzyme or preparations from untransfected cells; a positive control contained *Helix pomatia* sulfatase (Type H-1, Sigma-Aldrich, St. Louis, USA; 0.28 mg ml^−1^). Reactions were incubated at room temperature for 14 h or 2 h (depending on purpose) and stopped by adding 500 µl methanol, followed by cooling on ice or at −20 °C. After centrifugation (15,000 × g, 5–10 min, 4 °C), supernatants were dried under nitrogen and redissolved in water for UHPLC-HRMS (for product detection and identification) or HPLC-MS (for quantification of remaining substrate). To obtain differential activity profiles, enzyme concentrations were adjusted to prevent complete conversion of any glucosinolate within 2 h of incubation.

### GSS assay with *Hesperis matronalis* extract

Freeze-dried leaves of *H. matronalis* (20 mg) were boiled with 400 µl methanol for 4 min. After addition of 400 µl water and glass beads, extraction was continued in a paint shaker for 90 s and an ultrasonic bath for 4 min. The supernatant obtained after centrifugation was dried under nitrogen and redissolved in 1 ml water to obtain the plant extract. Plant extract (50 µl) was mixed with 5 µl 125 µM 4-pentenyl glucosinolate, 5 µl 126 µM benzyl glucosinolate, and 5-15 µl enzyme solution (depending on the specific GSS) in a final volume of 100 µl 50 mM Tris-HCl (pH 7.5). Negative controls were obtained by adding buffer (100 mM Tris-HCl, pH 7.5) instead of enzyme solution. After 2 h at room temperature, reactions were stopped by adding 500 µl methanol on ice. Supernatants obtained after centrifugation (15,000 g, 5-10 min, 4 °C) were dried under nitrogen and redissolved in 50 µl water for HPLC-MS.

### Arylsulfatase assay

The standard sulfatase substrate 4-methylumbelliferyl sulfate (80 µl of an 8.75 mM stock in 50 mM Tris-HCl, pH 7.5, 500 mM NaCl, 100 µM MgCl2, 100 µM MnCl2, 100 µM CaCl2) was incubated with 25 µl enzyme preparation and 95 µl of the same buffer at 37 °C for 2 h, following^13^. Negative controls used boiled enzyme or preparations from untransfected cells; a positive control contained *H. pomatia* sulfatase (Type H-1, Sigma-Aldrich, St. Louis, USA; 0.28 mg ml^−1^). Formation of 4-methylumbelliferone was monitored at 370 nm in one-minute intervals using a microplate reader and quantified from a calibration series of 4-methylumbelliferone (0.5–500 µM).

### UHPLC-HRMS

Samples were analyzed on a Vanquish Horizon UHPLC system equipped with a Hypersil Gold aQ column using a gradient of 0.1 % formic acid in water (A) and 0.1 % formic acid in methanol (B): 1.8 % B for 2 min, 1.8–100 % B over 14 min, and 100 % B for 3 min (0.25 ml min^−1^), injection volume 10 µl. The UHPLC was coupled to an Exploris 120 Orbitrap operated in full-scan mode (m/z 100–500; resolution 120 000) and monitoring the eluent between 2.5 and 13.0 min. Instrument settings were: positive H-ESI, 3400 V; sheath gas 3 arb; auxiliary gas 2 arb; purge gas 0 arb; ion-transfer-tube temperature 320 °C; RF lens 70 %. Data were processed in FreeStyle (v1.8). Incubations of the glucosinolate mixture with *H. pomatia* sulfatase were used to determine retention times and detection limits of desulfoglucosinolates (Extended Data Table 10).

### HPLC-MS of assays with glucosinolate mix

Samples were analyzed on an Agilent 1200 HPLC system equipped with a Hypersil Gold aQ column using a gradient of 0.1 % formic acid in water (A) and 0.1 % formic acid in methanol (B): 1.8 % B for 8 min, 1.8–100 % B over 12 min, and 100 % B for 2 min (0.25 ml min^−1^), injection volume 5 µl. The HPLC was coupled to a 3200 QTrap mass spectrometer operated in negative mode with multiple-reaction monitoring (MRM). Two transitions were monitored for each glucosinolate, the first of which was used for quantification (Extended Data Table 11). Quantification was based on peak areas and calibration with the glucosinolate mixture; matrix effects were ≤ 10 % for all glucosinolates. Mass-spectrometer settings were: curtain gas 40 psi; ionization voltage −4500 V; ion-source temperature 450 °C; ion-source gas 1 and gas 2 at 60 psi; collision gas medium; entrance potential −10 V; collision-cell exit potential −3 V. Data were processed in Analyst (v1.6.2).

### HPLC-MS of assays with *H. matronalis* extract

Samples were analyzed on an Agilent 1200 HPLC system equipped with a Hypersil Gold aQ column using a gradient of 0.1 % formic acid in water (A) and 0.1 % formic acid in methanol (B): 1.8 % B for 2 min, 1.8–100 % B over 14 min, and 100 % B for 3 min (0.25 ml min^−1^), injection volume 5 µl. The HPLC was coupled to a 3200 QTrap mass spectrometer operated in negative mode with precursor-ion scanning (m/z 310–600 in 0.5 s). Masses generating a fragment with m/z 97 were detected. Mass-spectrometer settings were: curtain gas 45 psi; ionization voltage −4500 V; ion-source temperature 400 °C; ion-source gas 1 at 35 psi and gas 2 at 50 psi; collision gas medium; declustering potential −55 V; entrance potential −10 V; collision energy −40 eV; collision-cell exit potential −3 V. Data were processed in Analyst (v1.6.2). Glucosinolate assignment followed^8^ based on retention times, precursor-ion m/z values, and the derived fragment at m/z 97. Conversion of glucosinolates was determined by comparing precursor-ion peak areas in enzyme assays with those in negative controls.

### Quantification and statistical analysis

Based on HPLC-MS quantification of glucosinolates in the enzyme reactions, the quantity of converted individual glucosinolates was determined for each enzyme, using enzyme preparations from three independent expressions of each GSS. Relative conversions were estimated by analysis of covariance, using the total amount of substrate converted across all glucosinolates in each assay as covariate. From these models, mean conversion estimates and 99 % confidence intervals were obtained for each substrate. Adjusted values were centered separately for GSS1 and GSS3 by subtracting the class-specific mean across all substrates, yielding relative substrate-preference profiles independent of overall activity level. Spearman correlations were computed from centered values. For GSS1 enzymes, a deviation index was calculated as the sum of absolute deviations from the substrate-specific mean.

### Static structural models

Protein structures were generated with AlphaFold3^62^. Because the web interface of AlphaFold3 does not support α-formylglycine (FGly), the catalytic FGly position was represented as glycine in all static models. Docking was performed using Protenix under Neurosnap (https://neurosnap.ai/), using mmseqs2 with 6 recycles, 200 sampling steps and 5 diffusion samples. Models were visualized with Mol* [63], ChimeraX^64^ or PyMOL^65^. Non-covalent protein-ligand interactions were evaluated with PLIP^66^.

### FGly parametrization

For molecular-dynamics simulations, the catalytic α-formylglycine (FGly) residue was indroduced by replacing the glycine placeholder and parametrizing the aldehyde-containing side chain with GAFF2, while backbone atoms used ff14SB types. Missing parameters were completed using parmchk2, and partial charges were obtained from ORCA^67^ RESP/ESP calculations. FGly was defined as a protein residue in tleap.

### Ligand parametrization and placement

The glucosinolate-derived ligand 1-methylethyl glucosinolate (1-me) was parametrized using antechamber (AM1-BCC charges, GAFF2 atom types). A mol2 file was completed with parmchk2, and the total charge was adjusted to −1.0. To ensure consistent starting geometries across all proteins, ten manually constructed ligand poses were generated. After structural alignment of the six GSS proteins in ChimeraX, 1-me was placed near the binding pocket at ∼2 nm distance from the catalytic Ca^2+^ ion. These pose-specific PDB files were loaded into tleap together with the ligand mol2 and frcmod files.

### System assembly

Protein, ligand, and Ca^2+^ ion were combined in tleap (AmberTools^68^) using ff14SB, GAFF2, TIP3P water, and the Amber ion library. Systems were neutralized, solvated with a 12 Å TIP3P buffer, and adjusted to an ionic strength of 0.2 M. Final topology and coordinate files were generated using saveamberparm.

### Molecular dynamics simulations

MD simulations were performed in OpenMM^69^ using PME electrostatics (1.0 nm cutoff, Ewald tolerance 1 x 10^−3^) and constraints on bonds involving hydrogen. A Langevin integrator (300 K, 1 ps^−1^ friction, 2 fs timestep) was used. After minimization, systems were equilibrated for 0.05 ns (NVT) and 0.2 ns (NPT with Monte Carlo barostat). A harmonic restraint between the ligand COM and Ca^2+^ was applied during equilibration and removed before production. Production simulations were run for 20 ns in the NPT ensemble without restraints.

### Force decomposition and analysis

A parallel OpenMM system was constructed to decompose nonbonded interactions into Lennard-Jones, Coulomb, and flat-bottom components, enabling per-group force analysis on ligand atoms.

### Adaptive trajectory sampling and reporting

Trajectory output frequency was adjusted dynamically based on Ca-sulfate COM distance and sulfate orientation (cos θ), using three sampling states (far, transition, bound). Frames were re-imaged by centering the protein and wrapping ligand coordinates. Every 500 steps, distances, energies, force magnitudes, axial force components, sulfate orientation, residue-71–Phe131 COM distance, and the Pro72–His198–Lys283 gate-triad angle were recorded.

## Code availability

The OpenMM script is profided in Extended Data 12.

## AI Assistance

Microsoft Copilot was used as an interactive tool to assist in learning and developing the parametrization of protein and ligand, as well as in writing molecular dynamics simulation scripts. All scripts were independently tested and validated through execution on local hardware. Microsoft Copilot and Mistral AI were used for language refinement.

## Data availability

Sequence data that support the findings of this study have been deposited in [repository] with the [primary] accession codes [accession codes].

## Acknowledgements

We thank our colleagues Peter Huemer, Wolfram Mey, Thierry Varenne, Paul Abram, and Yu Okamura for the generous provision of micromoth samples, Jean Keller for performing gene predictions, Ingo Ott and Ingo Rustenbeck for access to laboratory equipment, Ricardo C. Rodriguez de la Vega Cuellar for performing the molecular modelling runs, and Jacqui Shykoff for comments. We acknowledge the use of Microsoft Copilot for language refinement and editing of this manuscript. All scientific content, conclusions, and interpretations were generated and verified by the authors.

## Author contributions

J.K., H.V. and U.W. conceived the study and designed analyses. L.P., M.P., J.K. and H.V. collected insect specimens. M.D.K. and P.K. synthesized substrates. L.P., M.P. and H.V. performed experiments. J.K. developed the OTTO-TOTO test. J.K., L.P., M.P., H.V. and U.W. analyzed data. J.K. drafted the manuscript, with contributions from U.W., L.P. and H.V. All authors agreed to the final version of the manuscript.

## Additional information

Supplementary Information is available for this paper at [https://doi.org/10.1038/s41586-026-XXXX-y].

