## Supplemental Data for "Rapid Host Plant Adaptation via Ectopic Gene Conversion in Plutellid Moths"

### Extended Data Table 1. Glucosinolates combined in equimolar mixture for use as GSS substrates.

| Glucosinolate side chain | Glucosinolate abbreviation | Source | Purity [%] |
|---|---|---|---|
| 1-methylethyl | 1-me | synthesized[8] | 97.0 |
| 2-methylpropyl | 2-mp | | 98.0 |
| 1-methylpropyl | 1-mp | | 98.0 |
| 2-propenyl | 2-prop | Phytoplan, | 99.9 |
| 3-butenyl | 3-but | Heidelberg, | 98.1 |
| 4-pentenyl | 4-pent | Germany | 99.1 |
| 4-methylthiobutyl | 4-mtb | | 99.7 |
| 3-methylsulfinylpropyl | 3-msop | | 99.8 |
| 4-methylsulfinylbutyl | 4-msob | | 99.9 |
| 5-methylsulfinylpentyl | 5-msop | | 99.3 |
| 6-methylsulfinylhexyl | 6-msoh | | 96.5 |
| 8-methylsulfinyloctyl | 8-msoo | | 98.5 |
| 10-methylsulfinyldecyl | 10-msod | | 97.3 |
| 3-methylsulfonylpropyl | 3-msoop | | 99.7 |
| 2-phenylethyl | 2-pe | | 99.9 |
| benzyl | benzyl | | 99.2 |
| 4-hydroxybenzyl | pohb | | 99.9 |
| indol-3-ylmethyl | i3m | | 99.9 |
| 4-hydroxy-I3M | 4ohi3m | | 99.9 |
| 4-methoxy-I3M | 4moi3m | | 98.1 |
| 1-methoxy-I3M | 1moi3m | | 98.5 |

### Extended Data Table 2. Insect species with origin and source.

| Insect species | Origin | Source |
|---|---|---|
| *Plutella armoraciae* (Busck, 1912) | Canada | Paul Abram, Agassiz Research and Development Centre, Canada |
| *Plutella porrectella* (Linnaeus, 1758) | Canada | |
| *Eidophasia messingiella* (Fischer von Röslerstamm, 1840) | Austria | Peter Huemer, Naturwissenschaftliche Sammlung der Tiroler Landesmuseen, Austria |
| *Rhigognostis hufnagelii* (Zeller, 1839) | France | Thierry Varenne, Nice, France |
| *Leuroperna sera* (Meyrick, 1885) | Japan | Yu Okamura, Tokyo Metropolitan University, Japan |
| *Baerenschenkia umtrunkula* (Mey, 2011) | Namibia | Wolfram Mey, Leibniz Institute for Research on Evolution and Biodiversity, Germany |

### Extended Data Table 3. Sequencing technologies and genome assembly metrics.

| Species | Sequencing technology | Genome Size [Mb] | N50 | No. contigs | No. predicted genes | BUSCO Values |
|---|---|---|---|---|---|---|
| *Plutella armoraciae* | MinION, Illumina | 398,915 | 450,956 | 3,913 | 19,343 | C:98.5%[S:97.2%,D:1.3%],F:0.8%,M:0.7%,n:1367 |
| *Plutella porrectella* | MinION, Illumina | 455,390 | 855,451 | 1,088 | 19,297 | C:99.0%[S:93.7%,D:5.3%],F:0.4%,M:0.6%,n:1367 |
| *Eidophasia messingiella* | PacBio HiFi | 988,813 | 1,025,030 | 2,229 | 23,218 | C:98.0%[S:94.1%,D:3.9%],F:0.7%,M:1.3%,n:1367 |
| *Leuroperna sera* | MinION, Illumina | 348,870 | 430,549 | 1,338 | 21,807 | C:98.0%[S:95.4%,D:2.6%],F:0.7%,M:1.3%,n:1367 |
| *Rhigognostis hufnagelii* | PacBio HiFi | 534,261 | 515,364 | 2,813 | 19,677 | C:93.7%[S:90.3%,D:3.4%],F:2.8%,M:3.5%,n:1367 |
| *Baerenschenkia umtrunkula* | MinION, Illumina | 388,125 | 43,046 | 15,498 | n/a | C:81.8%[S:78.3%,D:3.5%],F:6.3%,M:11.9%,n:1367 |

### Extended Data Table 4. Gene copy numbers of GSS1 and GSS3.

| Insect species | No. *GSS1* copies | No. *GSS3* copies |
|---|---|---|
| *Plutella armoraciae* | 2 | 1 |
| *Plutella porrectella* | 2 | 1 |
| *Eidophasia messingiella* | 4 | 1 |
| *Leuroperna sera* | 2 | 1 |
| *Rhigognostis hufnagelii* | 1 | 2 |
| *Baerenschenkia umtrunkula* | 0 | 0 |
| *Plutella xylostella* | 2 | 1 |
| *Plutella australiana* | 1 | 1 |

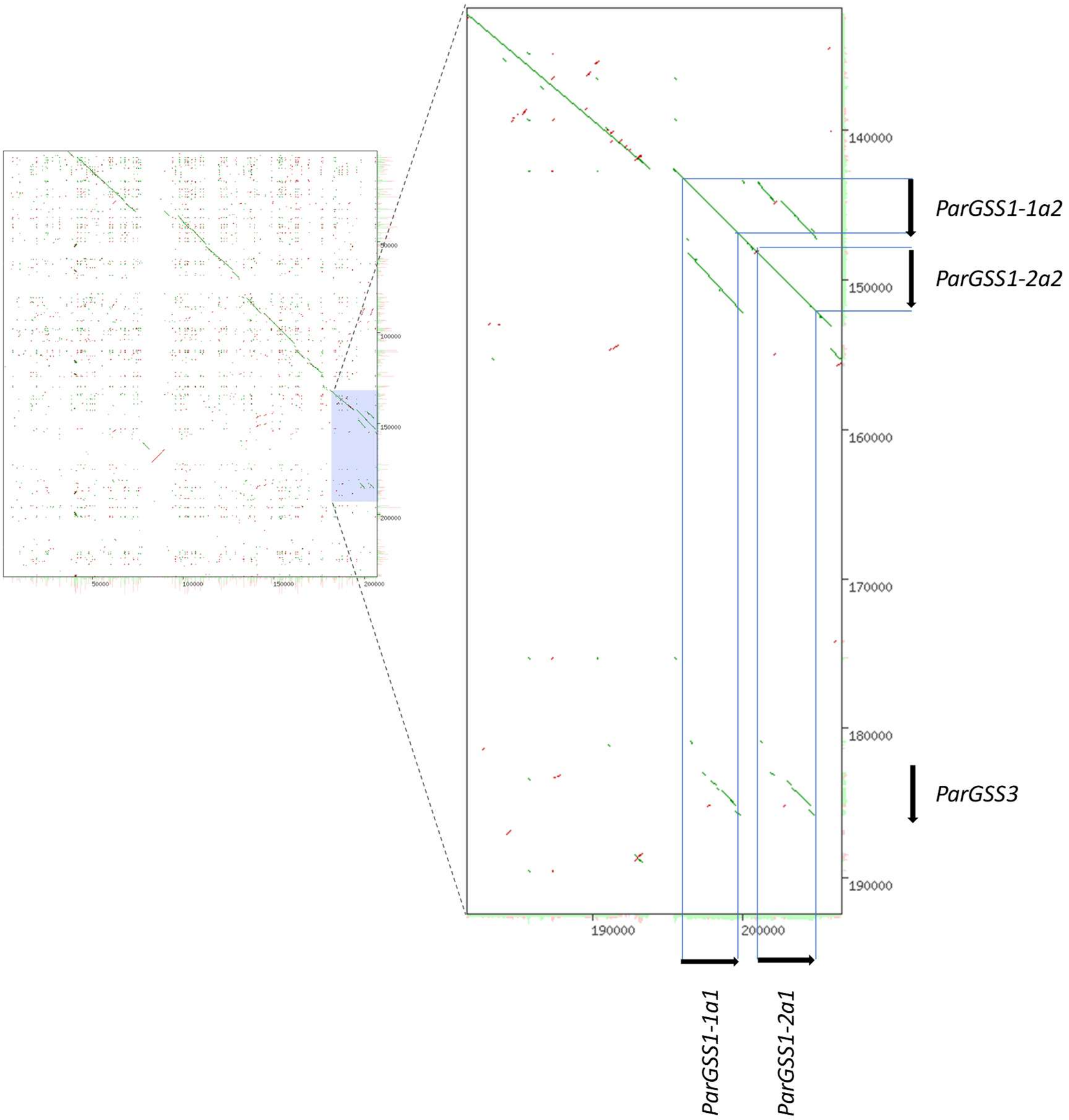


**Extended Data Fig. 1. Dot plot of allelic contigs containing *GSS* sequences in *Plutella armoraciae*.** The two contigs are co-linear (left), indicating that they represent alternative haplotypes rather than structural rearrangements. The magnified region on the right shows that the gene designated *ParGSS1-1a1* occupies the same genomic position as *ParGSS1-1a2*, demonstrating that these sequences are allelic variants of the same locus, even though ParGSS1-1a1 clusters with ParGSS1-2 in phylogenetic analyses.

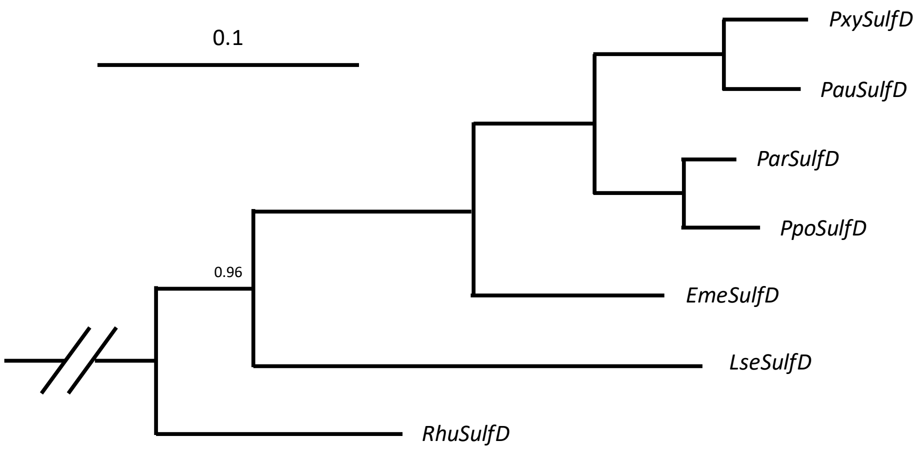


**Extended Data Fig. 2. *SulfD* gene tree.** The alignment spans codons 35 – 538 of the *P. xylostella SulfD* sequence, with *Yponomeuta cagnagella SulfD* as the outgroup. Scale bars indicate expected substitutions per codon site. All nodes show full posterior support except the single node annotated with a juxtaposed number. In *P. xylostella*, *SulfD* is tightly linked to the *GSS* gene cluster[13]. The inferred topology mirrors the species tree.

**Extended Data Table 5. Sequence identities between *GSS1* and *GSS3* in plutellid moths.**

| Species | Within-species identity [%] | Between-species identity [%] | Difference [percentage points] |
|---|---|---|---|
| *P. xylostella* | 68.95 | 67.61 | 1.3 |
| *P. australiana* | 70.50 | 67.68 | 2.8 |
| *P. armoraciae* | 69.95 | 68.18 | 1.8 |
| *P. porrectella* | 69.70 | 68.84 | 0.9 |
| *E. messingiella* | 70.38 | 68.09 | 2.3 |
| *L. sera* | 69.46 | 67.13 | 2.3 |
| *R. hufnaegelii* | 71.45 | 66.96 | 4.5 |

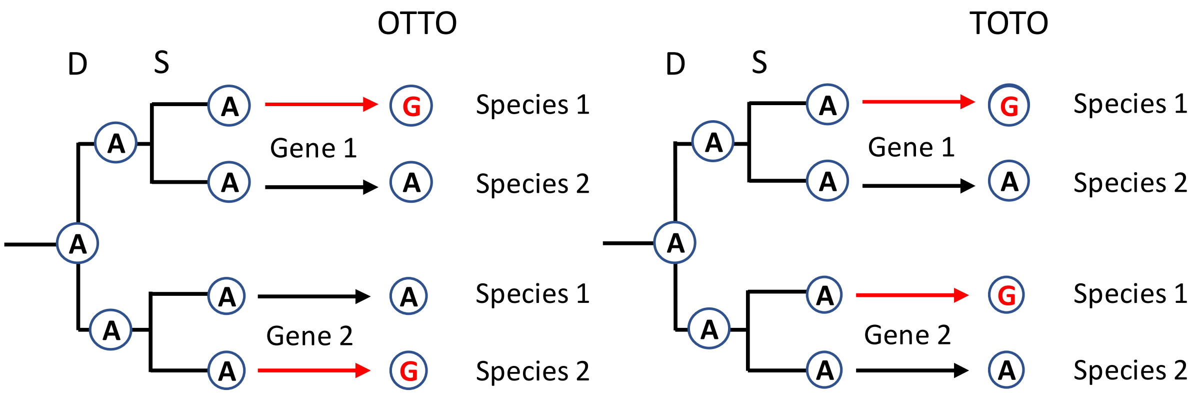


**Extended Data Fig. 3. Schematic representation of OTTO and TOTO configurations.** After a gene duplication (D) and a subsequent speciation (S) event, two independent point mutations (red arrows) occurring at the same site in the two paralogs can generate either an OTTO (left) or a TOTO (right) configuration with equal probability. In addition, a TOTO configuration can arise when an ectopic gene conversion (EGC) event homogenizes the two paralogs within a species. Because OTTO and TOTO should occur at similar frequencies under independent paralog evolution, a significant excess of TOTO over OTTO provides evidence for non-independent evolution driven by EGC.

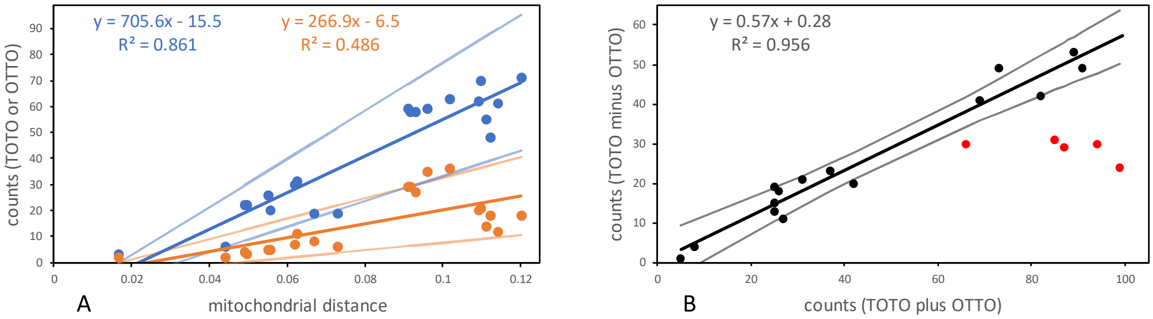


**Extended Data Fig. 4. Relationship between OTTO and TOTO configurations. (A)** Relationship between mitochondrial distance and the number of OTTO (orange) and TOTO (blue) configurations across all pairwise species comparisons. Shown are regression lines and 95% confidence intervals. Both OTTO and TOTO increased with distance (slopes: 266.9 vs. 705.6; 95 % CIs: [141.4 – 392.3] and [488.3 – 922.8]; *df* = 19). The slopes differed significantly (interaction term: $t \approx 4.9$; $df = 17$; $p < 0.001$), indicating faster accumulation of TOTO than OTTO configurations. Overall, TOTO counts were 2.7-fold (SE ± 0.24) more frequent than OTTOs (95% CI: 2.3 – 3.2; paired $t = 6.5$; $df = 20$; $p < 0.001$; $W = 231$; $p < 0.001$), revealing a pronounced excess beyond the neutral expectation of equal OTTO and TOTO counts. **(B)** Relationship between the absolute asymmetry (TOTO minus OTTO) and the total number of configurations (TOTO plus OTTO). Non-*Lse* pairs (red) followed a tight linear trend (slope 0.57; 95 % CI: [0.45 – 0.69]), whereas pairs involving *L. sera* consistently deviated downward, indicating a lineage-specific reduction in effective EGC, consistent with increased accumulation of independent point mutations following a major historical homogenization event.

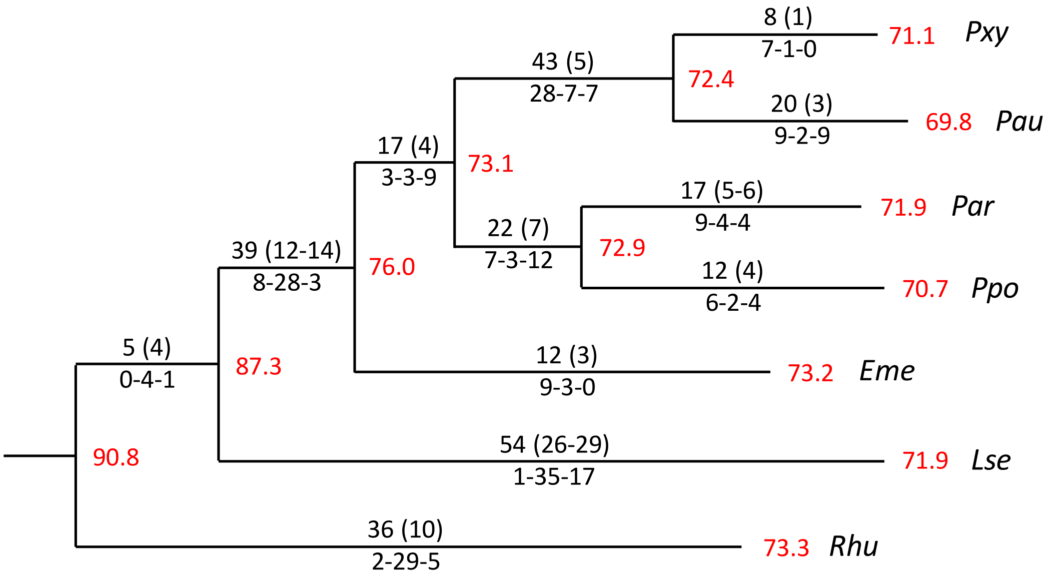


**Extended Data Fig. 5. Nucleotides putatively affected by EGC based on the OTTO-TOTO test, mapped onto the mitochondrial species phylogeny.** Numbers above branches indicate total and non-synonymous changes; numbers below branches indicate inferred directionality (*GSS1*→*GSS3*, undetermined, *GSS3*→*GSS1*). Red values next to nodes denote reconstructed pairwise identities of *GSS1* and *GSS3* coding sequences. For species with multiple *GSS1* or *GSS3* copies, the closest ancestral coding sequences were used for simplification. The OTTO-TOTO test identifies sites whose present state is compatible with an EGC event, but some TOTO configurations may instead reflect two independent point mutations, and earlier EGC events may have been obscured by subsequent substitutions. Estimates of nonsynonymous changes can be uncertain when later mutations altered neighboring nucleotides. Reconstructed ancestral identities are also influenced by EGC, which reduces divergence between *GSS1* and *GSS3*. All nucleotide positions identified as TOTO-compatible are listed in Extended Data Table 6.

**Extended Data Table 6. TOTO-compatible sites in the comparison of *GSS1* and *GSS3* coding sequences.**

| Species | Sites |
|---|---|
| *Pxy* | 129↓, 228↓, 660↓, 702↕, 1068↓, **1069↓**, 1515↓, 1593↓ |
| *Pau* | 153↕, 168↓, 180↕, 207↓, 345↓, 354↑, 375↑, 621↓, 696↓, 774↓, 858↑, 939↕, 981↓, **992↑**, **1078↑**, 1224↓, 1254↑, 1467↑, 1476↑, **1507↓**, 1644↑ |
| *Par* | 168↑, **331↓**, 459↓, 648↕, 657↕, 714↕, **1106↕**, 1158↓, 1176, **1223↓/1224↓**, 1404↓, 1467↓, 1494↓, **1498↑**, 1530↑, **1579↑** |
| *Ppo* | 660↕, 696↕, 705↓, 837↓, 960↑, 1179↓, **1204↑**, 1269↓, **1427↓**, **1507↑**, **1582↑**, 1638↓ |
| *Eme* | 139↓, **331↓**, 426↓, 606↓, 693↓, 696↓, 720↕, 739↓, **991↕**, **1325↓**, 1492↕, 1506↓ |
| *Lse* | **96↕**, 105↕, **139↕/141↕**, **142↑/143↑**, 180↕, 183↕, **250↑**, **320↑**, **346↑/347↕**, 426↑, **443↑**, 453↕, 591↕, **619↕/620↕**, 714↕, 739↑, 741↑, 742↑, 774↕, 858↕, **974↑**, **988↕**, **991↕**, 1029↓, **1078↑**, **1082↕**, **1123↕**, 1158↑, 1173↕, 1176↑, 1224↑, 1278↕, **1286↕/1287↕**, **1288↕**, **1291↑/1292↑**, **1321↕**, **1339↕/1340↕**, **1396↕**, 1401↕, 1404↕, **1405↕/1406↕**, **1436↕**, **1458↕**, **1464↕**, 1557↕, **1644↕** |
| *Rhu* | 73↕ 129↑, **142↓**, **147↕**, 153↕, **178↕**, 306↕, 348↕, 363↕, 366↑, 447↕, 450↕, 564↕, 738↕, 741↕, 744↕, **796↕**, **815↕/816↕**, 999↕, 1041↕, 1056↕, 1089↑, 1143↑, **1271↕**, **1308↕**, 1311↕, **1326↑**, 1392↕, **1427↕**, 1488↑, **1501↕**, 1521↕, 1560↕, 1569↕, 1578↕ |
| (*Pxy* , *Pau* ) | 153↓, 216↓, 315↕, **337↓**, **347↓**, **407↓**, 426↑, 459↓, 573↓, 603↓, 648↑, 669↓, 675↕, 693↕, 699↑, 705↓, 708↕, 742↓, 744↓, 753↓, 756↕, 795↑, **797↓**, 816↓, 825↓, 837↓, 888↓, 933↕, 960↓, 1008↓, **1021↓**, 1029↓, 1032↑, 1053↓, 1059↓, 1089↓, 1158↓, 1206↓, 1392↓, 1404↓, 1536↓, 1539↕, 1638↑ |
| (*Par* , *Ppo* ) | 81↑, 267↑, 309↓, 328↕, 387↕, 429↓, 453↑, 519↑, 648↓, 660↑, 696↑, 699↓, **799↕**, 888↑, **992↑**, 1032↓, 1053↑, **1216↓**, **1247↑**, **1325↑**, **1529↑**, **1606↓** |
| (*Pxy* , *Ppo* ) | **82↑**, 366↑, 534↓, 630↑, 693↑, 739↑, 756↕, **798↕**, 846↕, 939↕, **974↓**, 1071↕, 1278↑, 1320↑, 1326↑, 1458↑, **1488↓** |
| (*Pxy* , *Eme* ) | **74↓**, **145↕/147↕**, 345↕, 346↓, 354↕, 375↕, **463↑**, 486↕, **612↕**, 669↕, 681↕, 684↕, 705↕, 732↕, 741↓, **790↕**, **808↓**, 825↕, 933↕, 945↓, 948↕, **958↕**, 981↓, 1083↑, 1089↓, **1106↓**, 1143↑, 1254↕, **1432↕**, **1457↕**, 1476↕, 1539↕, 1542↕, 1558↕, **1568↕**, **1579↕/1581↕**, 1584↕ |
| (*Pxy* , *Rhu* ) | **715↕/716↕**, **736↕**, 774↑, **825↕** |

↓transfer from *GSS1* to *GSS3* ; ↕ undetermined; ↑ transfer from *GSS3* to *GSS1* ; orange: putative nonsynonymous change; red: nonsynonymous change; black: synonymous change

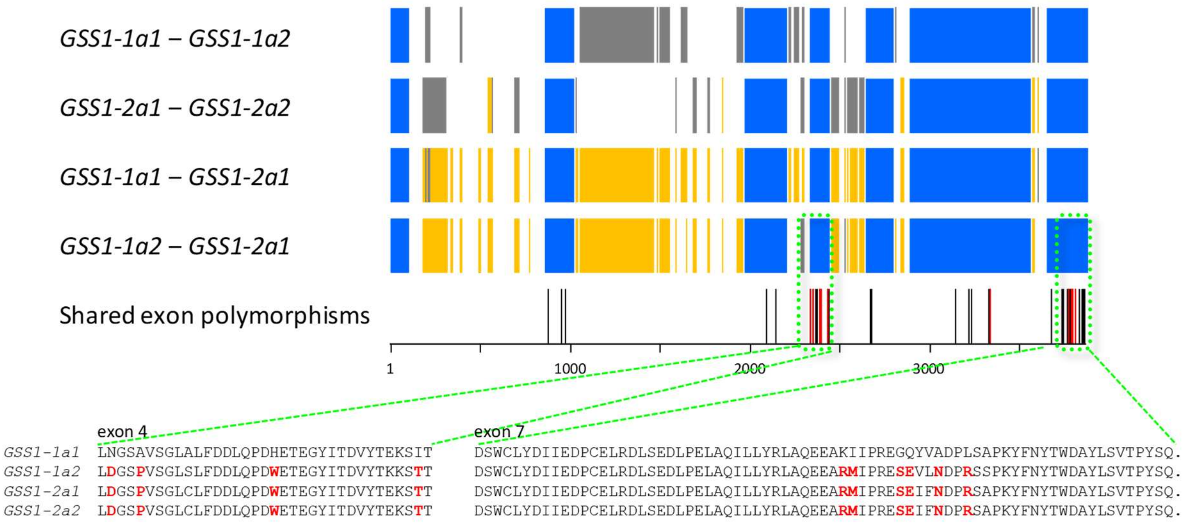


**Extended Data Fig. 6. Pairwise comparisons of genomic *ParGSS1* sequences.** Exons are shown in blue, gaps introduced during the multiple alignment in gray, and indel polymorphisms in orange. Synonymous (black) and non-synonymous (red) variants shared by *ParGSS1-1a2* and *ParGSS1-2*, but absent from *ParGSS1-1a1*, are highlighted. Below: amino-acid alignment of exons 4 and 7, with shared polymorphisms marked in red.

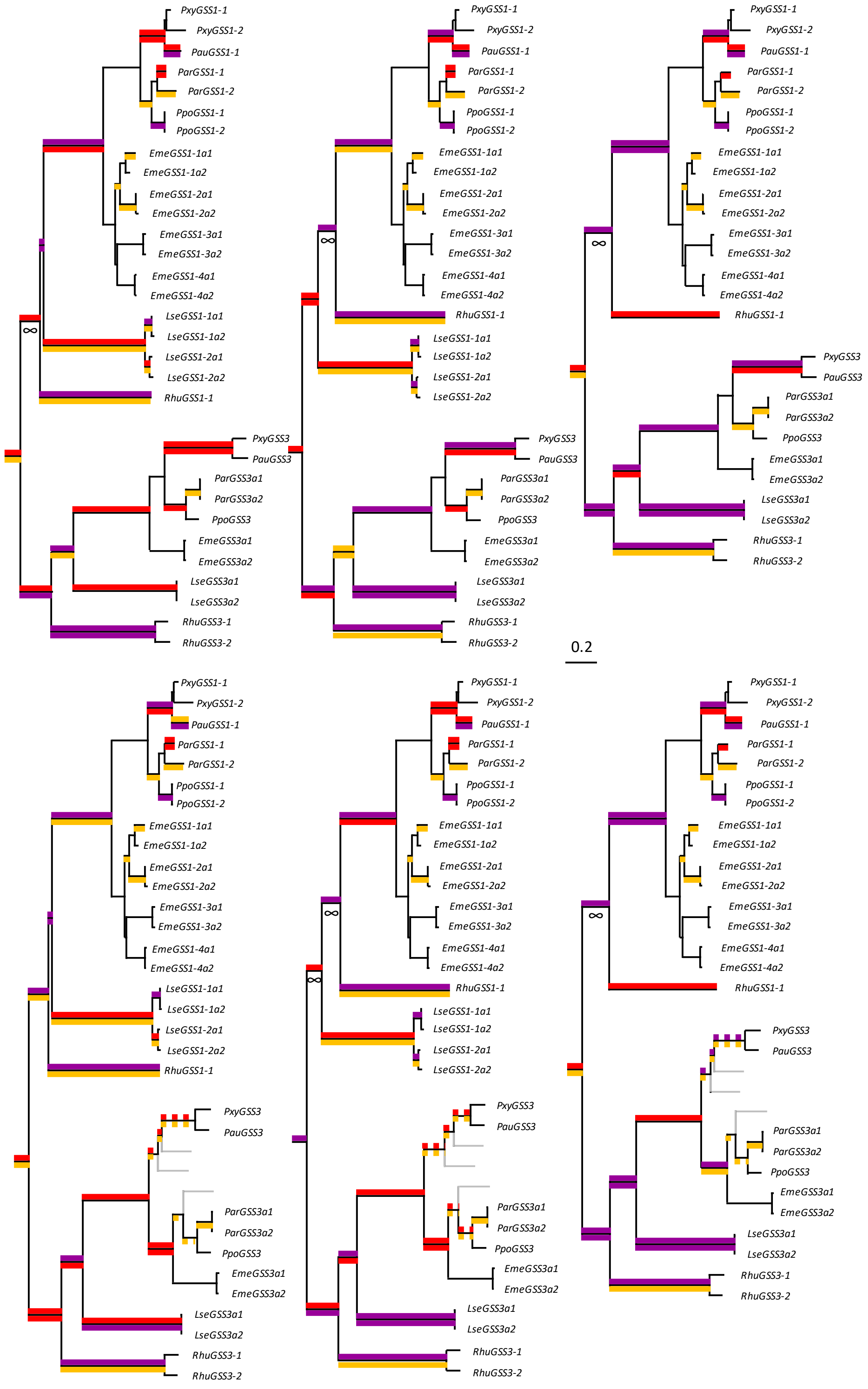


**Extended Data Fig. 7. Tests for positive selection.** We evaluated several alternative topologies by constraining branching orders in MrBayes – either involving *LseGSS1* and *RhuGSS1*, *EmeGSS3* vs.

*Plutella GSS3*, or by omitting *L. sera GSS1* sequences. The scale bar indicates substitutions per codon site. Significant signals in PAML branch-site models are shown above branches ($P_{adj}$ < 0.001 violet; $P_{adj}$ < 0.01 red; $P_{adj}$ < 0.05 orange; $P_{unadj}$ < 0.05 yellow), and aBSREL results below branches ($P_{adj}$ < 0.001 violet; $P_{unadj}$ < 0.01 red; $P_{unadj}$ < 0.05 orange). Across all tested topologies, both methods indicate strong positive selection. However, both assume substitutions arise independently through point mutations – an assumption violated by ectopic gene conversion.

**Extended Data Table 7. Plutellid GSS convert glucosinolates into desulfo-glucosinolates. Enzyme preparations were incubated with an equimolar glucosinolate mixture for 14 h followed by UHPLC-HRMS analysis.**

| Group | Identifier | 1-me | 2-mp | 1-mp | 2-prop | 3-but | 4-pent | 4-mtb | 3-msop | 4-msob | 5-msob | 6-msoh | 8-msoo | 10-msod | 3-msoop | 2-pe | benzyl | pohb | i3m | 4ohi3m | 4moi3m | 1moi3m |
|---|---|---|---|---|---|---|---|---|---|---|---|---|---|---|---|---|---|---|---|---|---|---|
| GSS1 | *Pxy* GSS1-1 | + | + | + | + | + | + | + | + | + | + | + | + | + | + | + | + | + | + | + | + | + |
| | *Pxy* GSS1-2 | + | + | + | + | + | + | + | + | + | + | + | + | + | + | + | - | + | + | + | - | - |
| | *Pau* GSS1-1 | + | + | + | + | + | + | + | + | + | + | + | + | + | + | + | + | + | + | + | + | + |
| | *Par* GSS1-1a1 | + | + | + | + | + | + | + | + | + | + | + | + | + | + | + | + | + | + | + | + | + |
| | *Par* GSS1-1a2 | + | + | + | + | + | + | + | + | + | + | + | + | + | + | + | + | + | + | + | + | + |
| | *Par* GSS1-2a1 | + | + | + | + | + | + | + | + | + | + | + | + | + | + | + | + | + | + | + | + | + |
| | *Ppo* GSS1-1 | + | + | + | + | + | + | + | + | + | + | + | + | + | + | + | + | + | + | + | + | + |
| | *Ppo* GSS1-2 | + | + | + | + | + | + | + | + | + | + | + | + | + | + | + | + | + | + | + | + | + |
| | *Eme* GSS1-1a1* | + | + | + | + | + | + | + | + | + | + | + | + | + | + | + | + | + | + | - | + | - |
| | *Eme* GSS1-1a2* | + | + | + | + | + | + | + | + | + | + | + | + | + | + | + | + | + | + | + | + | - |
| | *Eme* GSS1-2a2 | + | + | + | + | + | + | + | + | + | + | + | + | + | + | + | + | + | + | + | + | + |
| | *Eme* GSS1-3a2 | + | + | + | + | + | + | + | + | + | + | + | + | + | + | + | + | + | + | + | + | + |
| | *Eme* GSS1-4a1* | - | - | - | - | - | - | - | - | - | - | - | - | - | - | - | - | - | - | - | - | - |
| | *Lse* GSS1-1a1 | + | + | + | + | + | + | + | + | + | + | + | + | + | + | + | + | + | + | + | + | + |
| | *Lse* GSS1-1a2 | + | + | + | + | + | + | + | + | + | + | + | + | + | + | + | + | + | + | + | + | + |
| | *Lse* GSS1-2a1 | + | + | + | + | + | + | + | + | + | + | + | + | + | + | + | + | + | + | + | + | + |
| | *Lse* GSS1-2a2 | + | + | + | + | + | + | + | + | + | + | + | + | + | + | + | + | + | + | + | + | + |
| | *Rhu* GSS1 | + | + | + | + | + | + | + | + | + | + | + | + | + | + | + | + | + | + | - | + | - |
| GSS3 | *Pxy* GSS3 | + | + | + | - | + | + | + | - | + | - | - | + | + | - | + | + | + | + | + | + | + |
| | *Pau* GSS3 | + | + | + | + | + | + | + | + | + | + | + | + | + | + | + | + | + | + | + | + | + |
| | *Par* GSS3 | + | + | + | + | + | + | + | + | + | + | + | + | + | + | + | + | + | + | + | + | + |
| | *Ppo* GSS3 | + | + | + | + | + | + | + | + | + | + | + | + | + | + | + | + | + | + | + | + | + |
| | *Eme* GSS3 | + | + | + | + | + | + | + | + | + | + | + | + | + | + | + | + | + | + | + | + | + |
| | *Lse* GSS3 | + | + | + | + | + | + | + | + | + | + | + | + | + | + | + | + | + | + | + | + | + |
| | *Rhu* GSS3-1 | + | + | + | + | + | + | + | + | + | + | + | + | + | + | + | + | + | + | + | + | + |
| | *Rhu* GSS3-2 | + | + | + | + | + | + | + | + | + | + | + | + | + | + | + | + | + | + | + | + | + |
| Outgroup | *Pau* SulfD* | - | - | - | - | - | - | + | - | - | - | - | - | - | - | - | - | + | + | - | - | - |
| | *Rhu* SulfD* | - | - | - | - | - | - | - | - | - | - | - | - | - | - | - | - | - | - | - | - | - |
| | *Bombyx mori* ARS1* | - | - | - | - | - | - | + | - | + | - | - | + | + | - | + | + | + | + | + | + | + |
| Negative controls | untransfected cells | - | - | - | - | - | - | - | - | - | - | - | - | - | - | - | - | - | - | - | - | - |
| | untransfected cells | - | - | - | - | - | - | - | - | - | - | - | - | - | - | - | - | - | - | - | - | - |
| | untransfected cells | - | - | - | - | - | - | - | - | - | - | - | - | - | - | - | - | - | - | - | - | - |
| | boiled GSS | - | - | - | - | - | - | - | - | - | - | - | - | - | - | - | - | - | - | - | - | - |
| | boiled GSS | - | - | - | - | - | - | - | - | - | - | - | - | - | - | - | - | - | - | - | - | - |
| | boiled GSS | - | - | - | - | - | - | - | - | - | - | - | - | - | - | - | - | - | - | - | - | - |
| Positive controls | *Helix pomatia* sulfatase | + | + | + | + | + | + | + | + | + | + | + | + | + | + | + | + | + | + | + | + | + |
| | *Helix pomatia* sulfatase | + | + | + | + | + | + | + | + | + | + | + | + | + | + | + | + | + | + | + | + | + |
| | *Helix pomatia* sulfatase | + | + | + | + | + | + | + | + | + | + | + | + | + | + | + | + | + | + | + | + | + |

Enzyme preparations were incubated with an equimolar glucosinolate mixture for 14 h followed by UHPLC-HRMS analysis. '+ desulfo-glucosinolate detected; - desulfo-glucosinolate not detected; * no GSS signal in Western blots; all outgroups showed low product quantity

| Substrate (GSS1) | 1-me | 2-mp | 1-mp | 2-prop | 3-but | 4-pent | 4-mtb | 3-msop | 4-msob | 5-msop | 6-msoh | 8-msoo | 10-msod | 3-msoop | 2-pe | benzyl | pohb | i3m | 4ohi3m | 4moi3m | 1moi3m |
|---|---|---|---|---|---|---|---|---|---|---|---|---|---|---|---|---|---|---|---|---|---|
| 1-me | — | | | | | | | | | | | | | | | | | | | | |
| 2-mp | 0.34 | — | | | | | | | | | | | | | | | | | | | |
| 1-mp | 0.66 | 0.47 | — | | | | | | | | | | | | | | | | | | |
| 2-prop | 0.17 | -0.04 | 0.05 | — | | | | | | | | | | | | | | | | | |
| 3-but | -0.34 | 0.34 | 0.06 | 0.45 | — | | | | | | | | | | | | | | | | |
| 4-pent | -0.44 | 0.20 | -0.04 | 0.33 | 0.93 | — | | | | | | | | | | | | | | | |
| 4-mtb | -0.08 | 0.05 | -0.46 | 0.17 | 0.05 | 0.00 | — | | | | | | | | | | | | | | |
| 3-msop | 0.16 | -0.38 | -0.48 | 0.04 | -0.42 | -0.37 | 0.31 | — | | | | | | | | | | | | | |
| 4-msob | 0.03 | -0.72 | -0.44 | -0.04 | -0.58 | -0.55 | 0.28 | 0.51 | — | | | | | | | | | | | | |
| 5-msop | -0.23 | -0.42 | -0.51 | 0.01 | -0.35 | -0.32 | 0.42 | 0.23 | 0.72 | — | | | | | | | | | | | |
| 6-msoh | -0.48 | -0.09 | -0.40 | -0.09 | 0.08 | 0.15 | 0.15 | -0.11 | 0.19 | 0.57 | — | | | | | | | | | | |
| 8-msoo | -0.38 | 0.25 | -0.15 | -0.39 | 0.02 | 0.01 | 0.15 | -0.31 | -0.07 | 0.28 | 0.80 | — | | | | | | | | | |
| 10-msod | -0.40 | 0.09 | -0.19 | -0.42 | -0.06 | 0.03 | -0.05 | -0.30 | 0.00 | 0.28 | 0.80 | 0.91 | — | | | | | | | | |
| 3-msoop | -0.30 | -0.79 | -0.63 | 0.08 | -0.21 | -0.15 | 0.18 | 0.59 | 0.76 | 0.64 | 0.20 | -0.16 | -0.07 | — | | | | | | | |
| 2-pe | -0.21 | -0.02 | 0.01 | -0.29 | 0.15 | 0.14 | 0.26 | -0.04 | -0.04 | -0.34 | -0.08 | 0.20 | 0.07 | -0.20 | — | | | | | | |
| benzyl | 0.56 | 0.34 | 0.79 | 0.09 | 0.11 | 0.10 | -0.31 | -0.32 | -0.45 | -0.74 | -0.67 | -0.41 | -0.39 | -0.64 | 0.28 | — | | | | | |
| pohb | -0.09 | 0.04 | 0.28 | 0.16 | 0.37 | 0.33 | -0.47 | -0.10 | -0.58 | -0.57 | -0.46 | -0.45 | -0.45 | -0.12 | -0.09 | 0.26 | — | | | | |
| i3m | -0.18 | -0.56 | -0.32 | -0.30 | -0.63 | -0.60 | 0.06 | 0.09 | 0.58 | 0.35 | 0.02 | 0.02 | -0.01 | 0.29 | 0.06 | -0.26 | -0.31 | — | | | |
| 4ohi3m | -0.33 | -0.23 | -0.26 | -0.49 | -0.50 | -0.47 | -0.17 | -0.01 | 0.15 | 0.03 | -0.13 | 0.05 | 0.02 | -0.03 | 0.15 | -0.12 | -0.04 | 0.75 | — | | |
| 4moi3m | -0.22 | -0.19 | -0.09 | -0.69 | -0.58 | -0.50 | -0.28 | -0.08 | 0.13 | 0.24 | -0.05 | 0.05 | 0.11 | 0.07 | -0.15 | -0.20 | 0.09 | 0.53 | 0.74 | — | |
| 1moi3m | 0.10 | -0.41 | -0.29 | -0.33 | -0.73 | -0.74 | 0.11 | 0.38 | 0.56 | 0.17 | -0.29 | -0.22 | -0.25 | 0.25 | 0.06 | -0.13 | -0.25 | 0.87 | 0.71 | 0.47 | — |

| Substrate (GSS3) | 1-me | 2-mp | 1-mp | 2-prop | 3-but | 4-pent | 4-mtb | 3-msop | 4-msob | 5-msop | 6-msoh | 8-msoo | 10-msod | 3-msoop | 2-pe | benzyl | pohb | i3m | 4ohi3m | 4moi3m | 1moi3m |
|---|---|---|---|---|---|---|---|---|---|---|---|---|---|---|---|---|---|---|---|---|---|
| 1-me | — | | | | | | | | | | | | | | | | | | | | |
| 2-mp | 0.28 | — | | | | | | | | | | | | | | | | | | | |
| 1-mp | 0.60 | 0.74 | — | | | | | | | | | | | | | | | | | | |
| 2-prop | 0.74 | 0.67 | 0.74 | — | | | | | | | | | | | | | | | | | |
| 3-but | 0.66 | 0.07 | 0.03 | 0.18 | — | | | | | | | | | | | | | | | | |
| 4-pent | 0.85 | 0.40 | 0.54 | 0.66 | 0.72 | — | | | | | | | | | | | | | | | |
| 4-mtb | 0.31 | -0.36 | -0.17 | 0.00 | 0.38 | 0.40 | — | | | | | | | | | | | | | | |
| 3-msop | 0.70 | -0.07 | 0.41 | 0.61 | 0.11 | 0.49 | 0.58 | — | | | | | | | | | | | | | |
| 4-msob | 0.85 | 0.05 | 0.25 | 0.54 | 0.70 | 0.66 | 0.05 | 0.43 | — | | | | | | | | | | | | |
| 5-msop | 0.01 | -0.35 | -0.33 | -0.34 | 0.34 | 0.02 | 0.80 | 0.18 | -0.20 | — | | | | | | | | | | | |
| 6-msoh | 0.66 | 0.07 | 0.03 | 0.18 | 1.00 | 0.72 | 0.38 | 0.11 | 0.70 | 0.34 | — | | | | | | | | | | |
| 8-msoo | 0.74 | 0.31 | 0.54 | 0.39 | 0.60 | 0.47 | 0.13 | 0.36 | 0.57 | 0.27 | 0.60 | — | | | | | | | | | |
| 10-msod | 0.36 | 0.34 | 0.63 | 0.14 | 0.16 | 0.20 | 0.09 | 0.21 | 0.00 | 0.34 | 0.16 | 0.79 | — | | | | | | | | |
| 3-msoop | 0.39 | -0.30 | -0.15 | -0.07 | 0.52 | 0.01 | 0.20 | 0.20 | 0.51 | 0.46 | 0.52 | 0.72 | 0.38 | — | | | | | | | |
| 2-pe | -0.32 | 0.20 | 0.25 | -0.07 | -0.49 | -0.06 | 0.27 | 0.07 | -0.75 | 0.22 | -0.49 | -0.32 | 0.25 | -0.65 | — | | | | | | |
| benzyl | -0.74 | -0.45 | -0.74 | -0.46 | -0.49 | -0.53 | 0.11 | -0.29 | -0.57 | 0.04 | -0.49 | -0.93 | -0.79 | -0.49 | 0.25 | — | | | | | |
| pohb | -0.87 | 0.04 | -0.32 | -0.36 | -0.78 | -0.76 | -0.63 | -0.64 | -0.68 | -0.45 | -0.78 | -0.75 | -0.46 | -0.56 | 0.21 | 0.64 | — | | | | |
| i3m | -0.96 | -0.32 | -0.70 | -0.64 | -0.63 | -0.87 | -0.25 | -0.61 | -0.75 | 0.02 | -0.63 | -0.75 | -0.50 | -0.29 | 0.18 | 0.82 | 0.86 | — | | | |
| 4ohi3m | -0.63 | 0.22 | -0.20 | -0.32 | -0.32 | -0.38 | -0.76 | -0.86 | -0.36 | -0.61 | -0.32 | -0.57 | -0.43 | -0.56 | 0.00 | 0.36 | 0.79 | 0.54 | — | | |
| 4moi3m | -0.52 | -0.07 | 0.04 | -0.11 | -0.92 | -0.67 | -0.63 | -0.14 | -0.43 | -0.61 | -0.92 | -0.50 | -0.21 | -0.41 | 0.18 | 0.32 | 0.75 | 0.50 | 0.43 | — | |
| 1moi3m | -0.99 | -0.25 | -0.61 | -0.71 | -0.63 | -0.87 | -0.40 | -0.75 | -0.79 | -0.07 | -0.63 | -0.71 | -0.39 | -0.34 | 0.21 | 0.71 | 0.89 | 0.96 | 0.68 | 0.54 | — |

**Extended Data Fig. 8. Spearman correlation matrices of GSS1 (top) and GSS3 (bottom) substrate activities.** Colors indicate correlation strength (light red, moderate positive; red, strong positive; light blue, moderate negative; dark blue, strong negative). GSS1 (*df* = 13): $|\rho| \geq 0.514$ ($p \leq .05$); $|\rho| \geq 0.641$ ($p < .01$); $|\rho| \geq 0.770$ ($p < .001$). GSS3 (*df* = 5): $|\rho| \geq 0.714$ ($p \leq .05$); $|\rho| \geq 0.881$ ($p < .01$); $|\rho| \geq 0.952$ ($p < .001$).

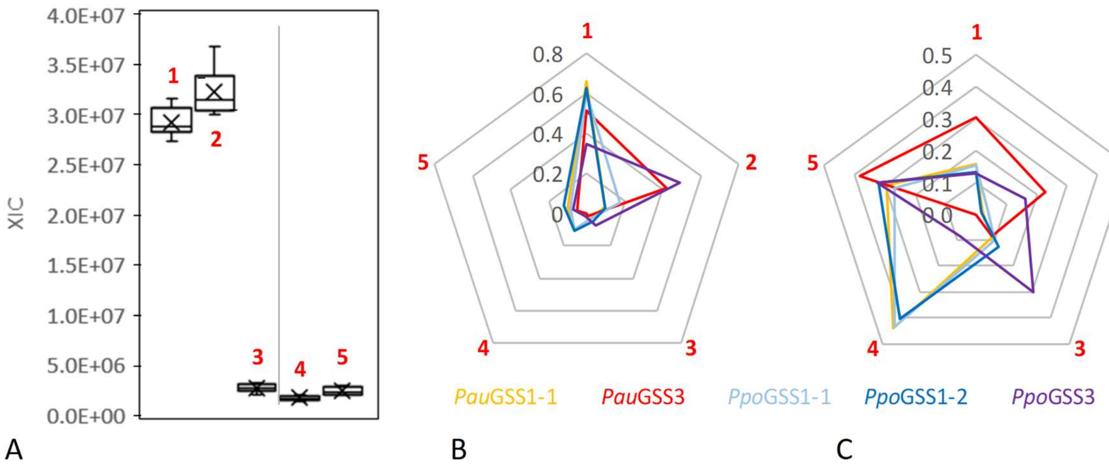


**Extended Data Fig. 9. Conversion of glucosinolates in *Hesperis matronalis* extract by GSS from *P. australiana* and *P. porrectella*.** (**A**) Major glucosinolates present in the reaction mixture (negative controls). Shown are peak areas from extracted-ion chromatograms (XIC) representing endogenous *H. matronalis* glucosinolates – 3,4-dihydroxy-benzyl (1, m/z 440), 4-apiosyloxy-3-hydroxybenzyl (2, m/z 572), and 4-apiosyloxybenzyl (3, m/z 556) – together with the added standard substrates 4-pentenyl (4, m/z 386) and benzyl (5, m/z 408). Peak areas from five negative control samples were measured twice (N=10) and are shown as box plots. (**B**) Normalized peak-area changes in enzyme reactions. For each substrate, the difference between mean peak areas in negative controls and individual enzyme reactions was normalized by the total peak-area change across all substrates within the same reaction. The resulting proportion of peak-area change attributable to GSS activity is shown as means from N = 3 independent expression experiments. (**C**) Concentration-corrected enzyme activity profiles. Normalized peak-area changes from panel (B) were divided by the corresponding substrate peak-area proportions in negative controls (panel A), yielding concentration-corrected relative activity. These values were then normalized by the total concentration-corrected peak-area change across all substrates for each enzyme.

All three GSS1 enzymes show similar substrate spectra but low activity toward apiosylated glucosinolates (2 and 3), whereas GSS3 from *P. porrectella* (for which *H. matronalis* is a primary host[29]) efficiently converts these unusual substrates. *Pau*GSS3 efficiently converts 4-apiosyloxy-3-hydroxybenzyl (2) glucosinolate but shows low activity toward 4-apiosyloxybenzyl glucosinolate (3), consistent with *P. australiana* being endemic to Australia where *H. matronalis* does not occur.

**Extended Data Table 8. Structural model quality metrics and confidence scores.**

| Ligand | Model ID | Overall Quality | Aggregate Score | Mean pLDDT | global PDE | pTM | ipTM | Uncertainty |
|---|---|---|---|---|---|---|---|---|
| 2-prop | *Par* GSS1-1a1 | High | 0.892±0.002 | 92.905±0.061 | 0.464±0.001 | 0.967±0.000 | 0.873±0.002 | 1.036±0.160 |
| | *Par* GSS1-1a2 | Very high | 0.948±0.000 | 94.011±0.018 | 0.383±0.001 | 0.972±0.000 | 0.942±0.000 | 0.662±0.113 |
| | *Par* GSS1-2 | Very high | 0.928±0.001 | 93.845±0.026 | 0.410±0.001 | 0.970±0.000 | 0.917±0.001 | 0.766±0.125 |
| | 1-1a1_e4 | Very high | 0.955±0.002 | 93.700±0.020 | 0.378±0.002 | 0.972±0.000 | 0.951±0.002 | 0.782±0.105 |
| | 1-1a1_e7 | Very high | 0.946±0.001 | 93.861±0.070 | 0.384±0.002 | 0.972±0.000 | 0.940±0.001 | 0.674±0.210 |
| | 1-1a1_e4e7 | Very high | 0.950±0.000 | 93.795±0.028 | 0.384±0.001 | 0.972±0.000 | 0.944±0.001 | 0.652±0.151 |
| 1-me | *Par* GSS1-1a1 | Very high | 0.907±0.003 | 93.744±0.028 | 0.438±0.003 | 0.970±0.000 | 0.892±0.004 | 0.816±0.083 |
| | *Par* GSS1-1a2 | Very high | 0.932±0.009 | 93.713±0.084 | 0.405±0.008 | 0.970±0.001 | 0.922±0.011 | 0.850±0.164 |
| | *Par* GSS1-2 | Very high | 0.907±0.001 | 93.343±0.078 | 0.436±0.002 | 0.968±0.000 | 0.891±0.002 | 1.148±0.232 |
| | 1-1a1_e4 | Very high | 0.957±0.002 | 93.754±0.027 | 0.377±0.002 | 0.972±0.000 | 0.953±0.003 | 0.916±0.089 |
| | 1-1a1_e7 | Very high | 0.958±0.002 | 93.594±0.047 | 0.378±0.002 | 0.972±0.000 | 0.954±0.002 | 0.902±0.080 |
| | 1-1a1_e4e7 | Very high | 0.958±0.001 | 92.481±0.035 | 0.396±0.002 | 0.970±0.000 | 0.955±0.001 | 1.096±0.092 |

Overall Quality: Overall structural quality assessment; Aggregate Score: Aggregate structural confidence score; Mean pLDDT: predicted Local Distance Difference Test (mean); global PDE: Predicted Distance Error (global); pTM: predicted Template Modeling score; ipTM: interface predicted Template Modeling score; Uncertainty: model ensemble uncertainty

**Extended Data Table 9. Residency times from MD simulations (sulfate-ester-to-Ca2+ distance ≤ 3.6 nm; ns).**

| Starting pose | *Par* GSS1-1a1 | *Par* GSS1-1a2 | *Par* GSS1-2 | 1-1a1_e4 | 1-1a1_e7 | 1-1a1_e4e7 |
|---|---|---|---|---|---|---|
| 1 | 20.0 | 20.0 | 18.5 | 20.0 | 20.0 | 20.0 |
| 2 | 20.0 | 20.0 | 20.0 | 20.0 | 20.0 | 9.2 |
| 3 | 20.0 | 1.9 | 20.0 | 5.0 | 19.5 | 9.5 |
| 4 | 11.6 | 20.0 | 20.0 | 20.0 | - | 11.1 |
| 5 | 20.0 | 12.1 | 12.8 | 15.2 | 12.5 | 20.0 |
| 6 | 8.6 | 3.0 | 19.7 | 4.4 | 20.0 | 7.6 |
| 7 | 3.8 | 20.0 | 2.9 | 20.0 | 18.9 | 18.9 |
| 8 | 14.7 | 6.0 | 20.0 | 20.0 | 20.0 | 20.0 |
| 9 | 12.5 | 6.1 | 20.0 | 5.9 | 0.5 | 8.9 |
| 10 | - | 2.8 | 20.0 | 20.0 | 20.0 | 19.7 |
| 1 | 11.6 | 20.0 | 20.0 | 8.9 | 20.0 | 20.0 |
| 2 | 20.0 | 5.8 | 20.0 | 8.9 | 19.3 | 16.8 |
| 3 | 20.0 | 20.0 | 20.0 | 20.0 | 20.0 | 20.0 |
| 4 | 3.2 | 20.0 | 20.0 | 19.6 | 7.5 | 20.0 |
| 5 | 20.0 | 20.0 | 20.0 | 20.0 | 1.9 | 6.2 |
| 6 | 20.0 | 20.0 | 9.8 | 6.2 | 20.0 | 20.0 |
| 7 | 20.0 | 20.0 | 20.0 | 20.0 | 16.4 | 20.0 |
| 8 | 20.0 | 11.3 | 15.8 | 9.4 | 20.0 | 20.0 |
| 9 | 20.0 | 3.4 | 19.7 | 8.1 | 20.0 | 20.0 |
| 10 | 4.6 | 16.3 | 20.0 | 20.0 | 13.8 | 19.9 |
| Mean | 15.3 | 13.4 | 18.0 | 14.6 | 16.3 | 16.4 |
| SEM | 6.3 | 7.5 | 4.5 | 6.5 | 6.3 | 5.3 |

**Extended Data Table 10. UHPLC-HRMS identification of desulfo-glucosinolates.**

| **Desulfo-glucosinolate (side chain)** | **m/z** | **tR** | **LOD [µM]** |
|---|---|---|---|
| 1-methylethyl | 282.10-282.10 | 6.33 | 0.10 |
| 2-methylpropyl | 296.11-296.12 | 7.50 | 0.10 |
| 1-methylpropyl | 296.11-296.12 | 7.82 | 0.10 |
| 2-propenyl | 280.08-280.09 | 5.16 | 0.10 |
| 3-butenyl | 294.10-294.10 | 6.92 | 0.01 |
| 4-pentenyl | 308.11-308.12 | 8.46 | 0.10 |
| 4-methylthiobutyl | 342.10-342.11 | 8.57 | 0.01 |
| 3-methylsulfinylpropyl | 344.08-344.08 | 3.56 | 0.10 |
| 4-methylsulfinylbutyl | 358.10-358.10 | 5.33 | 0.01 |
| 5-methylsulfinylpentyl | 372.11-372.12 | 6.32 | 0.01 |
| 6-methylsulfinylhexyl | 386.13-386.13 | 7.32 | 0.10 |
| 8-methylsulfinyloctyl | 414.16-414.16 | 9.52 | 0.01 |
| 10-methylsulfinyldecyl | 442.19-442.19 | 11.63 | 0.01 |
| 3-methylsulfonylpropyl | 360.08-360.08 | 3.81 | 0.10 |
| 2-phenylethyl | 344.11-344.12 | 9.85 | 0.01 |
| benzyl | 330.10-330.10 | 8.52 | 0.01 |
| 4-hydroxybenzyl | 346.09-346.10 | 6.70 | 0.01 |
| indol-3-ylmethyl | 369.11-369.11 | 9.02 | 0.01 |
| 4-hydroxyindol-3-ylmethyl | 385.10-385.11 | 7.37 | 0.01 |
| 4-methoxyindol-3-ylmethyl | 399.12-399.12 | 9.93 | 0.10 |
| 1-methoxyindol-3-ylmethyl | 399.12-399.12 | 10.90 | 0.01 |

Parameters were derived from a *Helix pomatia* sulfatase-treated glucosinolate mixture. tR, retention time; LOD, limit of detection

**Extended Data Table 11. Mass transitions and MS/MS parameters for detection and quantification of glucosinolates by HPLC-MS.**

| Glucosinolate (side chain) | Qualifier 1: Transition [Da] | DP [V] | CE [V] | Qualifier 2: Transition [Da] | DP [V] | CE [V] |
|---|---|---|---|---|---|---|
| 1-methylethyl | 360.0-97.0 | -55 | -30 | 360.0-75.0 | -55 | -50 |
| 2-methylpropyl | 374.0-97.0 | -58 | -30 | 374.0-75.0 | -58 | -55 |
| 1-methylpropyl | 374.0-97.0 | -58 | -30 | 374.0-75.0 | -58 | -55 |
| 2-propenyl | 358.0-97.0 | -43 | -28 | 358.0-75.0 | -43 | -55 |
| 3-butenyl | 372.0-97.0 | -60 | -30 | 372.0-75.0 | -60 | -50 |
| 4-pentenyl | 386.0-97.0 | -50 | -30 | 386.0-75.0 | -50 | -50 |
| 4-methylthiobutyl | 420.0-97.0 | -55 | -30 | 420.0-75.0 | -55 | -55 |
| 3-methylsulfinylpropyl | 422.0-97.0 | -55 | -35 | 422.0-358.0 | -60 | -30 |
| 4-methylsulfinylbutyl | 436.0-97.0 | -58 | -40 | 436.0-178.0 | -58 | -35 |
| 5-methylsulfinylpentyl | 450.0-97.0 | -56 | -40 | 450.0-386.0 | -56 | -30 |
| 6-methylsulfinylhexyl | 464.0-97.0 | -56 | -44 | 464.0-400.0 | -56 | -30 |
| 8-methylsulfinyloctyl | 492.0-97.0 | -50 | -50 | 492.0-75.0 | -50 | -50 |
| 10-methylsulfinyldecyl | 520.0-97.0 | -80 | -60 | 520.0-75.0 | -80 | -70 |
| 3-methylsulfonylpropyl | 438.0-97.0 | -55 | -40 | 438.0-75.0 | -55 | -55 |
| 2-phenylethyl | 422.0-97.0 | -55 | -35 | 422.0-75.0 | -50 | -50 |
| benzyl | 408.0-97.0 | -55 | -33 | 408.0-75.0 | -55 | -55 |
| 4-hydroxybenzyl | 424.0-97.0 | -54 | -35 | 424.0-75.0 | -54 | -55 |
| indol-3-ylmethyl I3M | 447.0-97.0 | -58 | -38 | 447.0-75.0 | -58 | -60 |
| 4-hydroxyindol-3-ylmethyl | 463.0-97.0 | -50 | -36 | 463.0-75.0 | -50 | -60 |
| 4-methoxyindol-3-ylmethyl | 477.0-97.0 | -55 | -39 | 477.0-75.0 | -60 | -60 |
| 1-methoxyindol-3-ylmethyl | 477.0-97.0 | -55 | -39 | 477.0-446.0 | -50 | -20 |

Qualifier 1 was used for quantification. DP, declustering potential; CE, collision energy

**Extended Data 12. OpenMM Script**

```
#!/usr/bin/env python

import os
import sys
import numpy as np
import glob

from openmm import *
from openmm.app import *
from openmm.unit import *

# ------------------------------------------------------------
# Hilfsfunktion für Dateiabfrage
# ------------------------------------------------------------

#def ask_file(prompt, default=None):
#    if default is not None:
#        s = input(f"{prompt} [{default}]: ").strip()
#        if s == "":
#            s = default
#    else:
#        s = input(f"{prompt}: ").strip()
#    if not os.path.isfile(s):
#        print(f"Datei nicht gefunden: {s}")
#        sys.exit(1)
#    return s

# Check if the correct number of arguments is provided
if len(sys.argv) != 3:
    print("Usage: python3 run_openmm_beam_v11R.py <stem>.prmtop <stem>.inpcrd")
    sys.exit(1)

# Assign the input files from command-line arguments
PRMTOP_FILE = sys.argv[1]
INPCRD_FILE = sys.argv[2]

# Extract the stem from the PRMTOP_FILE (e.g., "stem1" from "stem1.prmtop")
BASENAME = os.path.splitext(os.path.basename(PRMTOP_FILE))[0]

# Define output file paths
DCD_FILE = f"{BASENAME}.traj.dcd"
DIST_FILE = f"{BASENAME}.distance.txt"
STEPS_FILE = f"{BASENAME}.adaptive_steps.txt"

print(f">>> PRMTOP-Datei: {PRMTOP_FILE}")
```

```
print(f">>> INPCRD-Datei: {INPCRD_FILE}")
print(f">>> Trajektorie:   {DCD_FILE}")
print(f">>> Distance-Log:  {DIST_FILE}")
print(f">>> Steps-Log:     {STEPS_FILE}")

# Create a folder for this stem
os.makedirs(BASENAME, exist_ok=True)

# Update output file paths to include the stem folder
DCD_FILE = os.path.join(BASENAME, f"{BASENAME}.traj.dcd")
DIST_FILE = os.path.join(BASENAME, f"{BASENAME}.distance.txt")
STEPS_FILE = os.path.join(BASENAME, f"{BASENAME}.adaptive_steps.txt")

# ------------------------------------------------------------
# Auxiliary function for Beam
# ------------------------------------------------------------

def ask_yes_no(prompt, default="n"):
    default = default.lower()
    if default not in ("y", "n"):
        raise ValueError("Default muss 'y' oder 'n' sein")

    if default == "y":
        s = input(f"{prompt} [Y/n]: ").strip().lower()
        if s == "":
            return True
    else:
        s = input(f"{prompt} [y/N]: ").strip().lower()
        if s == "":
            return False

    return s.startswith("y")

# ------------------------------------------------------------
# Inputs
# ------------------------------------------------------------

#PRMTOP_FILE = ask_file("PRMTOP-Datei (.prmtop): ")
#INPCRD_FILE = ask_file("INPCRD-Datei (.inpcrd): ")


###I think this is already taken care of in the output block above###
#          BASENAME = os.path.splitext(os.path.basename(PRMTOP_FILE))[0]
#          DCD_FILE  = f"{BASENAME}.traj.dcd"
#          DIST_FILE = f"{BASENAME}.distance.txt"
#          STEPS_FILE = f"{BASENAME}.adaptive_steps.txt"
```

```
#
#           print(f">>> Trajectory:   {DCD_FILE}")
#           print(f">>> Distance-Log:  {DIST_FILE}")

# use_beam = ask_yes_no("Activate tractor beam?", default="n")
# lambda_value = 1.0 if use_beam else 0.0
lambda_value = 0.0 #falls Traktorstrahl gewünscht, die beiden vorherige Zeilen aktivieren und diese Zeile deaktivieren

###I think this will actually delete a file if already produced###
for f in [DCD_FILE, DIST_FILE, STEPS_FILE]:
    if os.path.exists(f):
        print(f"Delete old file: {f}")
        os.remove(f)
# Simulationszeiten
NVT_NS  = 0.05
NPT_NS  = 0.2
PROD_NS = 20.0

#making one tenth of the requested time
#           NVT_NS  = 0.005
#           NPT_NS  = 0.02
#           PROD_NS = 2.0


dt = 0.002 * picoseconds
steps_nvt  = int(NVT_NS  * 1000.0 / dt.value_in_unit(picoseconds))
steps_npt  = int(NPT_NS  * 1000.0 / dt.value_in_unit(picoseconds))
steps_prod = int(PROD_NS * 1000.0 / dt.value_in_unit(picoseconds))

print(f"NVT-Schritte:  {steps_nvt}")
print(f"NPT-Schritte:  {steps_npt}")
print(f"PROD-Schritte: {steps_prod}")

# ------------------------------------------------------------
# System aufbauen
# ------------------------------------------------------------

prmtop = AmberPrmtopFile(PRMTOP_FILE)
inpcrd = AmberInpcrdFile(INPCRD_FILE)

system = prmtop.createSystem(
    nonbondedMethod=PME,
    nonbondedCutoff=1.0*nanometer,
    constraints=HBonds
)
```

```
# >>> EwaldErrorTolerance im MD-System setzen <<<
for f in system.getForces():
    if isinstance(f, NonbondedForce):
        f.setEwaldErrorTolerance(1e-3)
        break

system.addForce(CMMotionRemover())

topology = prmtop.topology

# ------------------------------------------------------------
# Ca2+ und Sulfat-O-Atome finden
# ------------------------------------------------------------

ca_index = None
sulfate_O_indices = []
s_index = None

sulfate_O_names = {"O1", "O2", "O3"}

for atom in topology.atoms():
    if atom.element is not None and atom.element.symbol == "Ca":
        ca_index = atom.index

    if atom.residue.name == "1ME":
        if atom.name in sulfate_O_names:
            sulfate_O_indices.append(atom.index)
        if atom.name == "S":
            s_index = atom.index

if ca_index is None:
    print("Kein Ca2+ gefunden!")
    sys.exit(1)

if len(sulfate_O_indices) == 0:
    print("Keine Sulfat-O-Atome (O1/O2/O3) in 1ME gefunden!")
    sys.exit(1)

if s_index is None:
    print("Kein Schwefelatom (S) in 1ME gefunden!")
    sys.exit(1)

print(f"Ca-Index: {ca_index}")
print(f"Sulfat-O-Indizes: {sulfate_O_indices}")
print(f"S-Index: {s_index}")
```

```
# ----------------------------------------------------------
# Atomlisten für Residue 71 (Tyr oder Met) und Phe131
# ----------------------------------------------------------

top = topology   # Alias für Lesbarkeit

# --- Residue 131 bestimmen (PDB-Nummer 131 → index 130) ---
res131 = list(top.residues())[130]

phe131_ring_names = ["CG", "CD1", "CD2", "CE1", "CE2", "CZ"]

phe131_indices = [
    atom.index for atom in res131.atoms()
    if atom.name in phe131_ring_names
]

# --- Residue 71 bestimmen (PDB-Residuenummer 71 → index 70) ---
res71 = list(top.residues())[70]   # 0-basiert!

if res71.name == "TYR":
    res71_ring_names = ["CG", "CD1", "CD2", "CE1", "CE2", "CZ"]
    res71_indices = [
        atom.index for atom in res71.atoms()
        if atom.name in res71_ring_names
    ]

elif res71.name == "MET":
    res71_indices = [
        atom.index for atom in res71.atoms()
        if atom.name == "SD"
    ]

else:
    raise ValueError("Residue 71 ist weder TYR noch MET!")

print("Residue 71:", res71.name)
print("res71_indices:", res71_indices)
print("phe131_indices:", phe131_indices)

# ----------------------------------------------------------
# Gate-Triad: Pro72 – His198 – Lys283 (Winkel)
# ----------------------------------------------------------

res_pro72  = list(top.residues())[71]   # PDB 72 → index 71
res_his198 = list(top.residues())[197]  # PDB 198 → index 197
```

```
res_lys283 = list(top.residues())[282]  # PDB 283 → index 282

pro_atom_name = "CA"
his_atom_name = "CE1"   # robust gegen HIS/HIE/HID/HIP
lys_atom_name = "NZ"

pro72_idx = [a.index for a in res_pro72.atoms()  if a.name == pro_atom_name][0]
his198_idx = [a.index for a in res_his198.atoms() if a.name == his_atom_name][0]
lys283_idx = [a.index for a in res_lys283.atoms() if a.name == lys_atom_name][0]

print("Gate-Triad-Atome:")
print("Pro72 (CA):", pro72_idx)
print("His198 (CE1):", his198_idx)
print("Lys283 (NZ):", lys283_idx)

# ------------------------------------------------------------
# Ligand- und Protein-Atome bestimmen
# ------------------------------------------------------------

ligand_atoms = [a.index for a in topology.atoms() if a.residue.name == "1ME"]
# protein_atoms = [
#     a.index
#     for a in topology.atoms()
#     if a.residue.name not in ("1ME", "WAT", "HOH", "Na+", "Cl-")
# ]

# print("Ligand-Atome:", ligand_atoms[:10], "...")
# print("Protein-Atome:", len(protein_atoms))

masses_sulfate = np.array([
    system.getParticleMass(i).value_in_unit(dalton)
    for i in sulfate_O_indices
])

# ------------------------------------------------------------
# Analyse-System aufbauen
# ------------------------------------------------------------

analysis_system = prmtop.createSystem(
    nonbondedMethod=PME,
    nonbondedCutoff=1.0*nanometer,
    constraints=HBonds
)

# Original NonbondedForce im Analyse-System finden
```

```
orig_index_ana = None
for i, f in enumerate(analysis_system.getForces()):
    if isinstance(f, NonbondedForce):
        orig_index_ana = i
        break

orig_nb_ana = analysis_system.getForce(orig_index_ana)
orig_nb_ana.setEwaldErrorTolerance(1e-3)

# Neue Forces erzeugen
lj_force = NonbondedForce()
coul_force = NonbondedForce()

# Einstellungen kopieren
for nb_new in (lj_force, coul_force):
    nb_new.setNonbondedMethod(orig_nb_ana.getNonbondedMethod())
    nb_new.setCutoffDistance(orig_nb_ana.getCutoffDistance())
    nb_new.setUseSwitchingFunction(orig_nb_ana.getUseSwitchingFunction())
    nb_new.setSwitchingDistance(orig_nb_ana.getSwitchingDistance())
    nb_new.setUseDispersionCorrection(orig_nb_ana.getUseDispersionCorrection())
    nb_new.setEwaldErrorTolerance(orig_nb_ana.getEwaldErrorTolerance())

# Partikelparameter splitten
for i in range(orig_nb_ana.getNumParticles()):
    charge, sigma, epsilon = orig_nb_ana.getParticleParameters(i)
    lj_force.addParticle(0.0, sigma, epsilon)
    coul_force.addParticle(charge, sigma, 0.0)

# Exceptions splitten
for i in range(orig_nb_ana.getNumExceptions()):
    p1, p2, chargeProd, sigma, epsilon = orig_nb_ana.getExceptionParameters(i)
    lj_force.addException(p1, p2, 0.0, sigma, epsilon)
    coul_force.addException(p1, p2, chargeProd, sigma, 0.0)

# ForceGroups setzen
lj_force.setForceGroup(2)
coul_force.setForceGroup(3)

# Originale NB-Force entfernen
analysis_system.removeForce(orig_index_ana)

# Neue Forces hinzufügen
analysis_system.addForce(lj_force)
analysis_system.addForce(coul_force)

# Context erzeugen
```

```
analysis_integrator = VerletIntegrator(0.001*picoseconds)
analysis_context = Context(analysis_system, analysis_integrator)

# Boxvektoren übernehmen
if inpcrd.boxVectors is not None:
    analysis_context.setPeriodicBoxVectors(*inpcrd.boxVectors)

# ------------------------------------------------------------
# DistanceReporter
# ------------------------------------------------------------

class DistanceReporter(object):
    def __init__(self, filename, interval,
                 ca_index, s_index, sulfate_indices, ligand_atoms,
                 analysis_context, lj_group, coul_group, fb_group,
                 res71_indices, phe131_indices,
                 pro72_idx, his198_idx, lys283_idx,
                 masses):

        write_header = not os.path.exists(filename)

        self.filename = filename
        self.interval = interval
        self.ca_index = ca_index
        self.s_index = s_index
        self.sulfate_indices = sulfate_indices
        self.ligand_atoms = ligand_atoms
        self.res71_indices = res71_indices
        self.phe131_indices = phe131_indices
        self.pro72_idx = pro72_idx
        self.his198_idx = his198_idx
        self.lys283_idx = lys283_idx

        self.ctx = analysis_context
        self.lj_group = lj_group
        self.coul_group = coul_group
        self.fb_group = fb_group

        # Bitmask für alle drei ForceGroups
        self.force_mask = (1 << lj_group) | (1 << coul_group) | (1 << fb_group)

        if write_header:
            with open(self.filename, "w") as f:
                f.write(
                    "step\tdistance_nm\tpotential_kJmol\t"
                    "F_lj_mag\tF_coul_mag\tF_fb_mag\t"
```

```
            "F_lj_axis\tF_coul_axis\tF_fb_axis\torientation\tres71_phe131_nm\t"
            "angle_deg\n"
        )

    # Massen für COM
    self.masses = masses

def describeNextReport(self, simulation):
    steps = self.interval - simulation.currentStep % self.interval
    return (steps, True, False, False, True)

def report(self, simulation, state):

    positions = state.getPositions(asNumpy=True).value_in_unit(nanometer)

    # Positionen in Analyse-Context setzen
    self.ctx.setPositions(state.getPositions())

    ca_pos = positions[self.ca_index]
    sulfate_pos = positions[self.sulfate_indices]

    # Massen-gewichteter COM
    com = np.average(sulfate_pos, axis=0, weights=self.masses)

    # Distanz
    dist = np.linalg.norm(ca_pos - com)

    # Potentialenergie (MD-System)
    potE = state.getPotentialEnergy().value_in_unit(kilojoule_per_mole)

    # ------------------------------------------------------------
    # GPU-optimiert: EIN einziger getState()-Aufruf für ALLE Kräfte
    # ------------------------------------------------------------
    st = self.ctx.getState(getForces=True, groups=self.force_mask)
    forces = st.getForces(asNumpy=True).value_in_unit(kilojoule_per_mole/nanometer)

    # Kräfte für Ligand-Atome summieren
    F_total = forces[self.ligand_atoms].sum(axis=0)

    # Jetzt die Komponenten pro ForceGroup extrahieren
    # OpenMM liefert die Summe aller aktiven ForceGroups
    # → wir müssen die Beiträge einzeln holen
    # → dazu holen wir die Kräfte pro Gruppe separat aus dem Context
    #   aber OHNE neue Kraftberechnung (Context cached!)
    def get_group_force(group):
        stg = self.ctx.getState(getForces=True, groups=1 << group)
```

```
    fg = stg.getForces(asNumpy=True).value_in_unit(kilojoule_per_mole/nanometer)
    return fg[self.ligand_atoms].sum(axis=0)

F_lj   = get_group_force(self.lj_group)
F_coul = get_group_force(self.coul_group)
F_fb   = get_group_force(self.fb_group)

# Beträge
F_lj_mag = np.linalg.norm(F_lj)
F_coul_mag = np.linalg.norm(F_coul)
F_fb_mag = np.linalg.norm(F_fb)

# Achsenprojektion
r_vec = com - ca_pos
r_hat = r_vec / np.linalg.norm(r_vec)

F_lj_axis = np.dot(F_lj, r_hat)
F_coul_axis = np.dot(F_coul, r_hat)
F_fb_axis = np.dot(F_fb, r_hat)

# Orientierung
S_pos = positions[self.s_index]
v_SCa = ca_pos - S_pos
v_SCa_norm = v_SCa / np.linalg.norm(v_SCa)

cos_thetas = []
for O_idx in self.sulfate_indices:
    O_pos = positions[O_idx]
    v_SO = O_pos - S_pos
    v_SO_norm = v_SO / np.linalg.norm(v_SO)
    cos_thetas.append(np.dot(v_SCa_norm, v_SO_norm))

orientation = max(cos_thetas)

# Residue 71 – Phe131 Abstand
res71_com = positions[self.res71_indices].mean(axis=0)
phe131_com = positions[self.phe131_indices].mean(axis=0)
dist_71_131 = np.linalg.norm(res71_com - phe131_com)

# Gate-Triad Winkel
p = positions[self.pro72_idx]
l = positions[self.lys283_idx]
h = positions[self.his198_idx]
v1 = p - l
v2 = h - l
angle_deg = np.degrees(np.arccos(np.clip(
```

```
            np.dot(v1/np.linalg.norm(v1), v2/np.linalg.norm(v2)),
            -1.0, 1.0
        )))

        # Schreiben
        with open(self.filename, "a") as f:
            f.write(
                f"{simulation.currentStep}\t{dist:.3f}\t{potE:.1f}\t"
                f"{F_lj_mag:.1f}\t{F_coul_mag:.1f}\t{F_fb_mag:.1f}\t"
                f"{F_lj_axis:.1f}\t{F_coul_axis:.1f}\t{F_fb_axis:.1f}\t"
                f"{orientation:.3f}\t{dist_71_131:.3f}\t{angle_deg:8.1f}\n"
            )

# ------------------------------------------------------------
# AdaptiveDCDReporter (mit PBC-Reimaging)
# ------------------------------------------------------------

class AdaptiveDCDReporter:
    def __init__(self, filename, topology,
                 ca_index, s_index, sulfate_O_indices,
                 analysis_context,
                 steps_file=None,
                 far_interval=50000,
                 transition_interval=5000,
                 bound_interval=10000,
                 dist_transition=0.6,
                 dist_bound_enter=0.40,
                 ori_transition=0.6,
                 dist_bound_stable=0.35,
                 ori_bound=0.8,
                 stable_steps=5000,
                 report_interval=1000,
                 flush_interval=1):

        if steps_file is None:
            steps_file = f"{BASENAME}.adaptive_steps.txt"
#           steps_file = f"{BASENAME}.adaptive_steps.txt"

        self.steps_file = steps_file
        self.filename = filename
        self.topology = topology
        self.ca_index = ca_index
        self.s_index = s_index
        self.sulfate_O_indices = sulfate_O_indices
        self.ctx = analysis_context
```

```
        from openmm.app import DCDFile
        self.dcd = DCDFile(open(self.filename, "wb"), self.topology, 0)
        self.last_write = -999999
        self.frame_index = 0

        # State machine
        self.state = "far"
        self.stable_counter = 0

        # Parameters
        self.far_interval = far_interval
        self.transition_interval = transition_interval
        self.bound_interval = bound_interval
        self.dist_transition = dist_transition
        self.dist_bound_enter = dist_bound_enter
        self.dist_bound_stable = dist_bound_stable
        self.ori_transition = ori_transition
        self.ori_bound = ori_bound
        self.stable_steps = stable_steps
        self.report_interval = report_interval

        # Logfile öffnen
        self.steps_log = open(self.steps_file, "w")
        self.steps_log.write("frame\tstep\tstate\tdist\tori\n")
        self.steps_log.flush()

        # Precompute atom groups
        self.protein_indices = [i for i,a in enumerate(self.topology.atoms()) if a.residue.name != 'LIG']
        self.ligand_indices  = [i for i,a in enumerate(self.topology.atoms()) if a.residue.name == 'LIG']


    def describeNextReport(self, simulation):
        steps = self.report_interval - (simulation.currentStep % self.report_interval)
        return (steps, True, False, False, False)


    def report(self, simulation, state):

        # Positionen holen
        positions = state.getPositions(asNumpy=True).value_in_unit(nanometer)
        self.ctx.setPositions(state.getPositions())

        step = simulation.currentStep

        # Distanz & Orientierung
        ca_pos = positions[self.ca_index]
```

```
sulfate_pos = positions[self.sulfate_O_indices]
com = sulfate_pos.mean(axis=0)
dist = np.linalg.norm(ca_pos - com)

S_pos = positions[self.s_index]
v_SCa = ca_pos - S_pos
v_SCa_norm = v_SCa / np.linalg.norm(v_SCa)

cos_thetas = []
for O_idx in self.sulfate_O_indices:
    O_pos = positions[O_idx]
    v_SO = O_pos - S_pos
    v_SO_norm = v_SO / np.linalg.norm(v_SO)
    cos_thetas.append(np.dot(v_SCa_norm, v_SO_norm))

ori = max(cos_thetas)

# State machine
if self.state == "far":
    if dist < self.dist_transition and ori > self.ori_transition:
        self.state = "transition"
        self.stable_counter = 0

elif self.state == "transition":
    if dist < self.dist_bound_enter and ori > self.ori_bound:
        self.stable_counter += 1
        if dist < self.dist_bound_stable and self.stable_counter >= self.stable_steps:
            self.state = "bound"
    elif dist >= self.dist_transition:
        self.state = "far"

elif self.state == "bound":
    if dist > self.dist_bound_enter:
        self.state = "transition"
        self.stable_counter = 0

# Samplingrate
if self.state == "far":
    write_interval = self.far_interval
elif self.state == "transition":
    write_interval = self.transition_interval
else:
    write_interval = self.bound_interval

# Schreiben
if step - self.last_write >= write_interval:
```

```
            # --- PBC RE-IMAGING BLOCK ---
            state2 = simulation.context.getState(getPositions=True, enforcePeriodicBox=True)
            pos = state2.getPositions(asNumpy=True)
            box = state2.getPeriodicBoxVectors(asNumpy=True)

            # Protein zentrieren
            protein_com = np.mean(pos[self.protein_indices], axis=0)
            box_center = 0.5 * (box[0] + box[1] + box[2])
            shift = box_center - protein_com
            pos += shift

            # Ligand wrappen
            for i in self.ligand_indices:
                for dim in range(3):
                    L = box[dim][dim]
                    while pos[i][dim] < 0:
                        pos[i][dim] += L
                    while pos[i][dim] > L:
                        pos[i][dim] -= L

            self.dcd.writeModel(pos)

            self.last_write = step

            # adaptive_steps.txt
            self.steps_log.write(
                f"{self.frame_index}\t{step}\t{self.state}\t{dist:.4f}\t{ori:.4f}\n"
            )
            self.steps_log.flush()

            self.frame_index += 1

# ------------------------------------------------------------
# Flat-bottom corridor zwischen Ca2+ und Sulfat-COM
# ------------------------------------------------------------

fb = CustomCentroidBondForce(
    2,
    "lambda_fb * ( step(d - r_max) * k_fb * (d - r_max)^2"
    "            + step(r_min - d) * k_fb * (r_min - d)^2 );"
    "d = distance(g1, g2);"
)

# Globale Parameter
fb.addGlobalParameter("lambda_fb", 0.0)
```

```
fb.addGlobalParameter("r_min", 0.25 * nanometer)
fb.addGlobalParameter("r_max", 0.40 * nanometer)
fb.addGlobalParameter("k_fb", 1500.0 * kilojoule_per_mole/nanometer**2)

# Gruppe 0 = Ca2+
fb.addGroup([ca_index])

# Gruppe 1 = Sulfat-O-Atome (COM wird automatisch berechnet)
fb.addGroup(sulfate_O_indices)

# Bond zwischen Gruppe 0 und 1
fb.addBond([0, 1])

fb.setForceGroup(4)
fb_index = system.addForce(fb)

# ------------------------------------------------------------
# Integrator & Simulation
# ------------------------------------------------------------

integrator = LangevinIntegrator(300*kelvin, 1/picosecond, dt)
simulation = Simulation(topology, system, integrator)
# simulation.context.setParameter("k_rest", 0.0 * kilojoule_per_mole/nanometer**2)
simulation.context.setPositions(inpcrd.positions)

# ------------------------------------------------------------
# Startpositionen setzen
# ------------------------------------------------------------

# inpcrd.positions → numpy array
pos = np.array(inpcrd.positions.value_in_unit(nanometer))

# Ligand-COM bestimmen
lig_pos0 = pos[ligand_atoms].mean(axis=0)

# Boxvektoren setzen
if inpcrd.boxVectors is not None:
   simulation.context.setPeriodicBoxVectors(*inpcrd.boxVectors)

# ------------------------------------------------------------
# Relative Halteleine: Ligand-COM relativ zu Ca2+
# ------------------------------------------------------------

rel = CustomCentroidBondForce(
   2,
   "k_rel * (d - d0)^2; d = distance(g1, g2);"
```

```
)

rel.addGlobalParameter("k_rel", 300 * kilojoule_per_mole/nanometer**2)

# Startabstand bestimmen
lig_pos0 = pos[ligand_atoms].mean(axis=0)
ca_pos0 = pos[ca_index]
d0 = np.linalg.norm(lig_pos0 - ca_pos0)
rel.addGlobalParameter("d0", d0)

# Gruppen definieren
rel.addGroup(ligand_atoms)   # Ligand-COM
rel.addGroup([ca_index])     # Ca2+

rel_index = system.addForce(rel)

# ------------------------------------------------------------
# Pocket automatisch erkennen
# ------------------------------------------------------------

# r_cut = 0.8  # nm
# pocket_atoms = [
#     i for i in protein_atoms
#     if np.linalg.norm(pos[i] - lig_pos0) < r_cut
# ]

# print("Automatisch erkannte Pocket-Atome:", pocket_atoms)

# ------------------------------------------------------------
# (A) Repulsive Wall Ligand <-> Pocket
# ------------------------------------------------------------

# wall = CustomNonbondedForce("step(r0 - r) * k_rep * (r0 - r)^2")
# wall.addGlobalParameter("r0", 0.5*nanometer)
# wall.addGlobalParameter("k_rep", 2000*kilojoule_per_mole/nanometer**2)

# for _ in range(system.getNumParticles()):
#     wall.addParticle(())

# Exclusions aus LJ-Force übernehmen, damit alle Forces identische Paare haben
# for i in range(lj_force.getNumExceptions()):
#     p1, p2, _, _, _ = lj_force.getExceptionParameters(i)
#     wall.addExclusion(p1, p2)

# wall.addInteractionGroup(set(ligand_atoms), set(pocket_atoms))
# wall.setNonbondedMethod(CustomNonbondedForce.CutoffPeriodic)
```

```
# wall.setCutoffDistance(1.0*nanometer)

# wall_index = system.addForce(wall)

# ------------------------------------------------------------
# (B) COM-Leine um Startposition
# ------------------------------------------------------------

print("Ligand-Atome:", ligand_atoms)
print("Anzahl Ligand-Atome:", len(ligand_atoms))

# ------------------------------------------------------------
# COM-Leine als ExternalForce (robust, reinitialize-sicher)
# ------------------------------------------------------------

# Ligand-COM bestimmen
# state0 = simulation.context.getState(getPositions=True)
# pos0 = state0.getPositions(asNumpy=True)
# lig_pos0 = pos0[ligand_atoms].mean(axis=0)

# Federkonstante pro Atom
# k_total = 50 * kilojoule_per_mole / nanometer**2
# k_atom = k_total / len(ligand_atoms)

# hold = CustomExternalForce("k_atom * ((x-x0)^2 + (y-y0)^2 + (z-z0)^2)")
# hold.addGlobalParameter("k_atom", k_atom)
# hold.addGlobalParameter("x0", lig_pos0[0])
# hold.addGlobalParameter("y0", lig_pos0[1])
# hold.addGlobalParameter("z0", lig_pos0[2])

# for idx in ligand_atoms:
#     hold.addParticle(idx, [])

# hold_index = system.addForce(hold)

# Boxvektoren setzen
# if inpcrd.boxVectors is not None:
#     simulation.context.setPeriodicBoxVectors(*inpcrd.boxVectors)


# ------------------------------------------------------------
# Barostat (for NPT + simulation)
# ------------------------------------------------------------

barostat = MonteCarloBarostat(1*bar, 300*kelvin, 25)
```

```
# ------------------------------------------------------------
# Minimierung
# ------------------------------------------------------------

print(">>> Minimierung")
simulation.minimizeEnergy(maxIterations=2000)

# ------------------------------------------------------------
# NVT
# ------------------------------------------------------------

print(">>> NVT Equilibration")
simulation.context.setVelocitiesToTemperature(300*kelvin)
if steps_nvt > 0:
    simulation.step(steps_nvt)

# ------------------------------------------------------------
# NPT (with barostat)
# ------------------------------------------------------------

print(">>> NPT Equilibration (with barostat)")
system.addForce(barostat)
simulation.context.reinitialize(preserveState=True)
simulation.context.setVelocitiesToTemperature(300*kelvin)
simulation.step(steps_npt)

# ------------------------------------------------------------
# Remove restraints
# ------------------------------------------------------------

# print(">>> Entferne Wall + COM-Leine vor Produktion")

# Beide Forces in der richtigen Reihenfolge entfernen
# for idx in sorted([wall_index, hold_index], reverse=True):
#     system.removeForce(idx)

# simulation.context.reinitialize(preserveState=True)

# simulation.context.setParameter("k_rest", 100.0 * kilojoule_per_mole/nanometer**2)

# ------------------------------------------------------------
# Entfernen der relativen Halteleine
# ------------------------------------------------------------

print(">>> Entferne relative Halteleine vor Produktion")
system.removeForce(rel_index)
```

```
simulation.context.reinitialize(preserveState=True)
simulation.context.setVelocitiesToTemperature(300*kelvin)

# -----------------------------------------------------------
# Traktorstrahl (sanftes Einschalten)
# -----------------------------------------------------------

if lambda_value > 0.0:
    print("Scotty, beam me up...")
    n_steps = 100  # Anzahl der Ramp-Schritte
    for i in range(n_steps):
        lam = (i + 1) / n_steps
        simulation.context.setParameter("lambda_fb", lam)
        simulation.step(10)  # 10 Schritte pro Ramp-Level
        simulation.context.setVelocitiesToTemperature(300*kelvin)
    print("Traktorstrahl ist jetzt vollständig aktiv (lambda_fb = 1.0).")
else:
    print("Traktorstrahl bleibt deaktiviert (lambda_fb = 0.0).")

# -----------------------------------------------------------
# Simulation
# -----------------------------------------------------------

print(">>> Sampling")

# Reporter-Liste leeren
simulation.reporters.clear()

adaptive_dcd = AdaptiveDCDReporter(
    DCD_FILE, simulation.topology,
    ca_index, s_index, sulfate_O_indices,
    analysis_context
)
simulation.reporters.append(adaptive_dcd)

dist = DistanceReporter(
    DIST_FILE, 500,
    ca_index, s_index, sulfate_O_indices, ligand_atoms,
    analysis_context, lj_force.getForceGroup(), coul_force.getForceGroup(), fb.getForceGroup(),
    res71_indices, phe131_indices,
    pro72_idx, his198_idx, lys283_idx, masses_sulfate
)
simulation.reporters.append(dist)

simulation.step(5000)
```

```
simulation.step(steps_prod)

for r in simulation.reporters:
    if isinstance(r, AdaptiveDCDReporter):
        r.steps_log.flush()
        r.steps_log.close()

print("Done.")
```
